# Encapsulation-enhanced genetic switches in lactobacilli

**DOI:** 10.1101/2025.01.10.632477

**Authors:** Marc Blanch-Asensio, Varun Sai Tadimarri, Roberto Martinez, Gurvinder Singh Dahiya, Cao Nguyen Duong, Rahmi Lale, Shrikrishnan Sankaran

## Abstract

*Lactiplantibacillus plantarum* is known for its potential in healthcare, food production, and environmental biotechnology. However, its broader utility is constrained by a limited genetic toolbox, particularly lacking robust genetic switches for inducible gene expression. Addressing this gap, we developed a novel genetic switch for *L. plantarum* based on a strong bacteriophage-derived promoter and the food-grade inducer, cumate. However, the switch was susceptible to leaky expression in the late log phase of bacterial growth, which was correlated to a reduction in the culture pH. This leakiness was partially resolved by regulating culture conditions (temperature and nutrients) to limit growth below a certain bacterial density. More interestingly, leaky expression could be stably suppressed by encapsulating the bacteria in alginate as an engineered living material. This physically restricted growth and limited the pHdrop, thereby enhancing the switch performance. The possibilities to regulate protein secretion over several days, reversibly switch protein production, and establish dual functionalities by co-encapsulating strains with different switches were demonstrated. Thus, for the first time, we show a material-based strategy to enhance the performance of a genetic switch in bacteria. This strategy facilitates the development of *L. plantarum* for advanced applications in biotechnology, pharmaceutics, and living therapeutics.

## INTRODUCTION

Lactobacilli are highly valued for their diverse applications across healthcare, food production, and environmental remediation [1–3]. In addition, lactobacilli are natural inhabitants of multiple parts of the human body [4]. For instance, lactobacilli are the most frequently isolated bacteria from the normal vaginal microbiota and are involved in maintaining vaginal health [5]. Among their notable advantages is their ability to modulate gut microbiota, lower cholesterol, and attenuate inflammatory diseases [6, 7]. These characteristics highlight their potential as microbial chassis in diverse applications [8]. Building on their inherent benefits, there has been growing interest in harnessing lactobacilli through genetic engineering for advanced biotechnological applications with the highest number of reports based on the species *Lactiplantibacillus plantarum* [9]. For example, engineered lactobacilli have been modified to produce recombinant proteins for pharmaceutical applications, including vaccines and mucosal therapeutics [10, 11]. Recently, we demonstrated the unique benefits of encapsulating *L. plantarum* in alginate to create engineered living materials (ELMs) as a controlled drug-release platform [12].

However, the applicability of lactobacilli as living therapeutics or in ELMs is hindered by their limited genetic toolbox. In the last decades, progress has been made concerning finding natural or engineered promoters to drive high levels of gene expression in *L. plantarum* [13–15]. Nevertheless, lactobacilli still lack high-performance genetic parts like strong promoters, effective repressors, and reliable genetic switches for inducible gene expression compared to model bacteria such as *E. coli* [9]. This gap has slowed the development of advanced genetic circuits needed for more precise control of gene expression. Notably, reliable genetic switches have been particularly elusive to create. A few studies report expression systems such as xylose-, lactose-, and galactose-responsive genetic switches but the fold changes upon induction are relatively poor in these reported cases (<13x) [16, 17]. Bile-induced promoters have also been explored in lactobacilli, but their induction relies on intestinal conditions, which restrict their applicability [18]. The most reliable and widely reported genetic switch in *L. plantarum* is the pSIP system, which has been used to control the production of a wide range of proteins, including different enzymes and antigens [19–23]. Nevertheless, this switch is based on a two-component system limiting its adaptability, drives only moderate levels of expression, and relies on a peptide as an inducer, making it relatively cost-intensive.

Genetic switches are vital for the genetic programmability of an organism since they allow temporal control over gene expression [24, 25]. In terms of recombinant protein production for biotechnology or pharmaceutic applications, this enables a separation between growth and production phases, which can enhance productivity and reduce metabolic burden [26]. For living therapeutic or bioremediation applications, genetic switches enable biosensing functions or remote activation of desired responses like protein secretion [27]. Such switches are also key components for the generation of smart living materials able to sense and respond to the external environment [28]. However, inducible genetic switches in lactobacilli are plagued by limitations such as leaky expression, low expression levels on induction, and limited dynamic ranges, for which underlying reasons and reliable solutions are not known or reported in the literature.

Our recent studies have expanded the genetic programmability of *L. plantarum* by developing a high-performance promoter-repressor system derived from bacteriophages with the potential to aid the development of reliable genetic switches [29]. Moreover, towards the applicability of such genetically engineered lactobacilli, we encapsulated them in core-shell alginate hydrogels to create the Protein Eluting Alginate with Recombinant Lactobacilli (PEARL) format [12]. These ELMs provided benefits such as containment of engineered bacteria, sustained protein release, and reduced metabolic toxicity for at least two weeks. Moreover, it stabilized the protein release profile of the engineered bacteria when compared to non-encapsulated bacterial cultures. These results suggested that the PEARL format influenced the metabolism of the encapsulated bacteria and could probably enhance the performance of engineered *L. plantarum*.

Thus, in this study, we build upon these advances to create a novel genetic switch for *L. plantarum* that responds to a food-grade inducer molecule, cumate [24]. We uncover an interesting correlation between bacterial growth leading to a drop in the pH and the leakiness of the genetic switch. More interestingly, by encapsulating these engineered strains in the PEARL format, we discovered that the bacteria remain trapped in a state that favorably minimizes leaky expression and maintains high levels of switchable protein production even up to nine days. We further demonstrate the possibility of extending this approach to another highly leaky genetic switch, induced by vanillate, and leverage this capability to develop dual-switchable PEARLs. Finally, we show that this leakiness-suppressing-effect is sustained even when the PEARLs are miniaturized by an order of magnitude. These interesting results suggest that this encapsulation strategy maximizes the performance of genetic switches in *L. plantarum,* and this ELM approach could be promising for further development of gene regulation in lactobacilli.

## RESULTS AND DISCUSSION

### Developing a genetic switch in *L. plantarum*

To create a genetic switch in *L. plantarum*, we leveraged our previously discovered phage-derived strong constitute promoter, *P_tec_*. We modified the promoter with operator sequences associated with three switchable repressors from promising food-grade small-molecule inducible gene expression systems that were previously established in other bacteria - the anhydrotetracycline (aTc) system, proven to work in multiple bacteria such as *Escherichia coli*, *Vibrio cholerae* or *Streptococcus suis* [30–32], the cumate system, which functioned well in *Pseudomonas aeruginosa* and *Bacillus* subtilis [33, 34], and the vanillate system, which showed promising results in *E.* coli [35]. Accordingly, the *P_tec_* promoter was flanked by the corresponding operator sequences, one placed upstream and the other downstream of the promoter sequence (**Figure S1A** and **S1B**).

All three switches were designed with the same plasmid architecture, which harbored a *P_256_* origin of replication and erythromycin resistance gene. The mCherry reporter gene was positioned downstream of the operator-flanked *P_tec_* promoter, while the repressor for each genetic switch was placed under the control of a moderate-strength constitutive promoter, *P_23_* [13] (**Figure S1A**). This was done to ensure that the repressor gene was continuously expressed, while mCherry expression could be switched ON and OFF upon the addition of the respective inducers (aTc, cumate, and vanillate) (**Figure 1A**). Using flow cytometry, we assessed the responsiveness of each genetic switch to the given inducer after 5 hours of induction, seeking a candidate that strongly represses *P_tec_* in the absence of the inducer and triggers strong protein production upon induction (**Figure 1B**). The aTc switch (*P_23_*_tetR_mCherry variant) showed low leaky mCherry expression in the absence of the inducer but also very weak levels of expression upon induction with 0.2 - 4 µM of aTc (∼3x fold change vs uninduced, **Figure 1C** and **S1C**). The cumate switch (*P_23_*_cymR_mCherry variant) was more promising, with low levels of leakiness when uninduced (similar to wild-type *L. plantarum* WCFS1) and significantly higher levels of mCherry expression upon induction with 100 µM of cumate (∼27x fold change vs uninduced, **Figure 1C**). Finally, the vanillate switch (*P_23_*_vanR_mCherry variant) showed elevated levels of leakiness when uninduced and only slightly higher levels of expression with 50 - 200 µM vanillate (∼2x fold change vs uninduced, **Figure 1C** and **S1D**). Overall, all three switches were responsive to their inducers, but since the cumate switch offered the best performance, we proceeded to further characterize and improve this switch.

**Figure 1:**
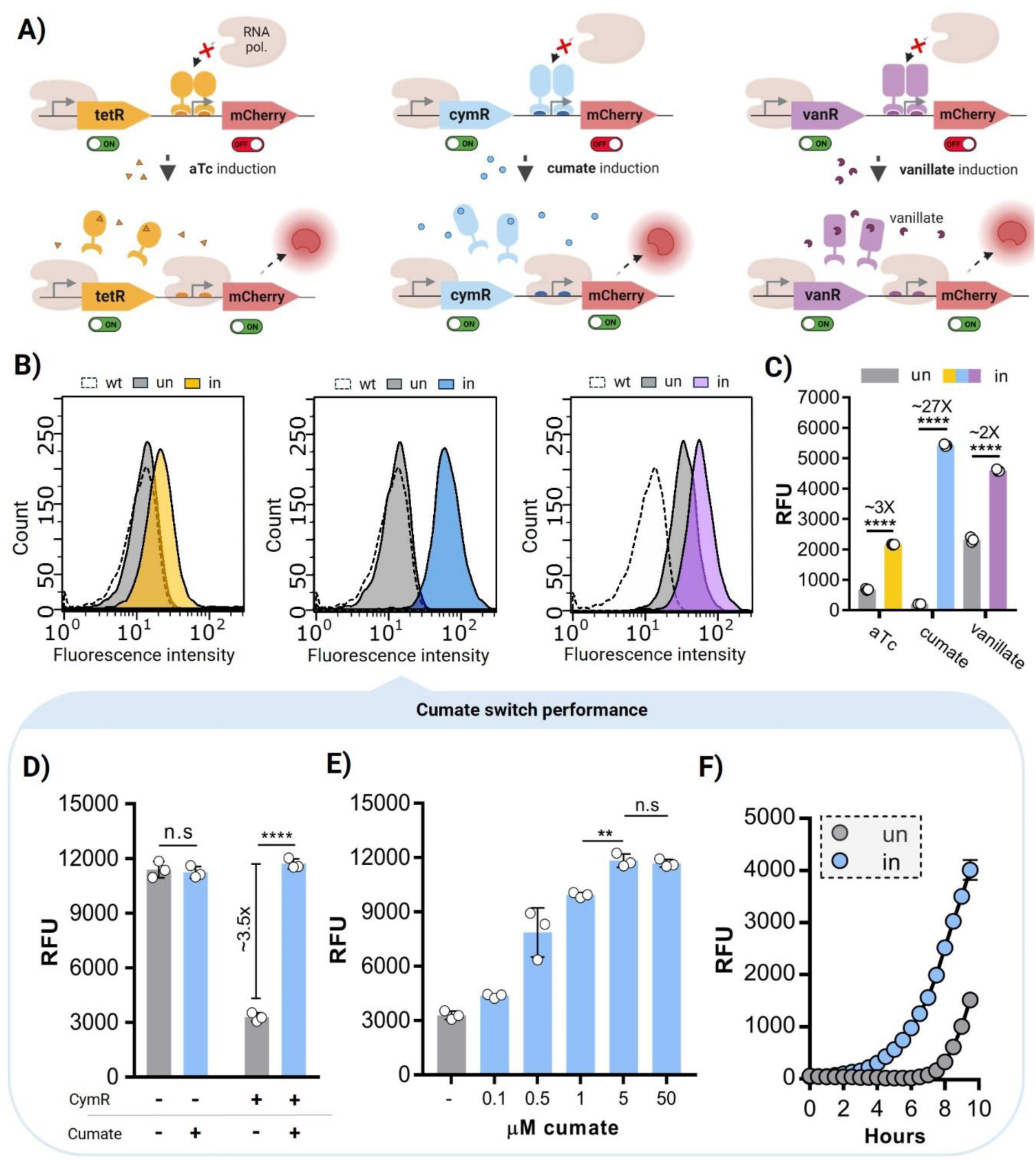
**A)** Schemes showing the mode of action of the three inducible switches, and the expected outcome after the addition of the inducers. **B)** Histograms obtained by flow cytometry showing the red fluorescence intensity of the uninduced and induced states of each switch. For all, the fluorescence intensity of non-engineered *L. plantarum* is shown as reference for leakiness. Bacteria were subcultured to an OD_600nm_ of 0.05 and induced for 6 hours with 0.2 µM of aTc, 100 µM of cumate and 50 µM of vanillate, respectively. **C**) RFU values for the aTc, cumate and vanillate switches after 5 hours of induction (starting OD_600nm_ = 0.05, n = 3, mean ± SD). “un” = uninduced condition, “in” = induced condition (0.2 µM of aTc, 100 µM of cumate and 50 µM of vanillate). **D**) RFU values for the *P_23_*_cymR_mCherry variant after 8-h induction with 100 µM of cumate. The presence of the repressor cymR in the plasmid and the addition of cumate is shown as “+”. The absence of the repressor cymR and cumate is shown as “-“ (starting OD_600nm_ = 0.05, n = 3, mean ± SD). **E)** RFU values for the *P_23_*_cymR_mCherry after inducing with different cumate concentrations for 8 hours (starting OD_600nm_ = 0.05, n = 3, mean ± SD). **F**) RFU values for the cumate switch after inducing with 5 µM of cumate for 10 hours (starting OD_600nm_ = 0.01, n = 3, mean ± SD). “un” = uninduced condition, “in” = induced condition (5 µM of cumate).

For that, we first tested whether the genetic switch responds to cumate only due to inhibition of the CymR repressor. Hence, in the plasmid encoding this genetic switch, we removed the *cymR* gene and observed that bacteria constitutively produced as much mCherry as the variants with the repressor that were induced by cumate (100 µM). This not only proved that induction was mediated by the repressor but also suggested that cumate fully inactivated it (**Figure 1D**). By varying the inducer concentration, we further determined that 5 µM cumate was sufficient for maximal induction, below which expression levels dropped in a seemingly linear manner. This suggested the possibility to tune expression levels by varying the concentration of cumate in the culture (**Figure 1E**).

Nevertheless, we observed considerable leakiness in the switch, indicating that repression of the inducible gene is incomplete in the absence of cumate. Notably, leakiness seemed to increase with longer culture times as indicated by the different fold changes at 5 h (**Figure S1C**) and 8 h (**Figure 1C**) after induction. We speculated that this difference may be associated with bacterial density in the cultures. So, we studied the performance of the cumate switch over time with cultures having different starting bacterial densities - OD_600nm_ of 0.3, 0.1, and 0.01 and correlated the data with kinetic measurements of bacterial growth. Interestingly, regardless of the initial bacterial density, leaky expression was observed in the uninduced cultures once bacteria crossed a density of 0.4 OD_600nm_ (measured in a microplate reader)(**Figure S2A** and **S2C**). This was in the late log-phase of growth, indicating that the operated *P_tec_* promoter is repressed during the early and mid-log phases of growth (**Figure S2B**).

We speculated that the leakiness could be associated with insufficient expression of the CymR repressor driven by the *P_23_* promoter. Hence, we developed two additional variants by replacing *P_23_* with *P_tlpA,_* a stronger constitutive promoter (*P_tlpA_*_cymR_mCherry variant), and with the natural (non-operated) *P_tec_* promoter (*P_tec_*_cymR_mCherry variant) (**Figure S3A**). Surprisingly, we observed similar trends as with the *P_23_*_cymR_mCherry variant (**Figure 1E**), with leakiness occurring in the uninduced state after 6-8 hours of growth (**Figure S3B** and **S3C**). Prominently, all variants supported similar levels and kinetics of protein production on induction. This suggested that failure of the repressor was not due to its quantity in the cell, but some other factor associated with growth.

Since building a genetic switch based on two independent genes (the repressor gene and the inducible gene) did not address the leakiness problem, we redesigned the switch employing a feedback circuit where both the repressor (cymR) and gene of interest (mCherry) are under the same operated *P_tec_* promoter, hypothetically achieving gene expression only in the presence of cumate (**Figure S3D**). This feedback mechanism has been used in genetic switches in *E. coli* to improve the stability of the switch, but it has never been attempted in *L. plantarum* [36, 37]. To ensure the production of both proteins, we included the *P_23_* ribosome-binding site between the STOP codon of the mCherry and the START codon of the cymR gene (**Figure S3D**). We termed this variant *P_tec_*_cymR_P_mCherry (polycistronic). This variant showed similar results concerning the leakiness of the switch, however, interestingly, the overall levels of production were higher (**Figure S3E**).

When comparing all these constructs in terms of induced fold change over uninduced leaky expression, the *P_23_*_cymR_mCherry switch emerges as the superior system with a maximal fold change above 60x at 6 hours (**Figure S3F**). However, it is to be noted that fold changes for all switches vary over time being low at that start before induced expression is detectable, increasing to a maximum while leaky expression remains poor, and reducing again as leaky expression increases.

Next, we compared the performance of *P_23_*_cymR_mCherry (best fold change switch) and *P_tec_*_cymR_P_mCherry (highest production switch) with the most widely used genetic switch in *L. plantarum*, the pSIP system. This is a two-component system in which an inducer peptide (SppIP) binds a transmembrane receptor kinase, which in turn phosphorylates a response regulator that acts as an activator to initiate transcription from the *P_spp_* promoter [20]. As expected, we observed clear differences between the performance of the cumate and the pSIP switches. Firstly, the pSIP switch showed negligible leakiness at all the assessed bacterial densities (0.3, 0.1, and 0.01). However, the overall levels of expression were significantly lower compared to the cumate switches (**Figure S4A**). For instance, after 12 hours of induction, mCherry production by the pSIP_409 system, was 8 to 17 times lower than that of the cumate switches (**Figure S4B**). Thus, the cumate switches based on the strong *P_tec_* promoter outperformed the pSIP switch in terms of protein production but were also leakier. Nevertheless, these results showed that it is possible to repress the operated *P_tec_* promoter during the early and mid-log phases of growth. Finding a way to decrease the leakiness of the cumate switch and increase the uninduced-induced state fold change would add great value to the genetic programmability of *L. plantarum*.

### Improving the cumate switch performance

From previous experience, we were aware that the strength of many promoters in *L. plantarum* are temperature-dependent [38]. Similarly, we observed that the *P_tec_* promoter is two-fold stronger at 37°C compared to 30°C (**Figure S5A**). Furthermore, temperature also affects the growth rate of bacteria with typically slower growth at lower temperatures. Based on these, we speculated that temperature could influence the performance of the cumate genetic switch by offering lower leakiness at 30°C and strong induced protein production at 37°C. We termed this concept as “*thermally aided switching*”, involving both temperature and cumate to regulate the performance of the switch (**Figure 2A**).

**Figure 2:**
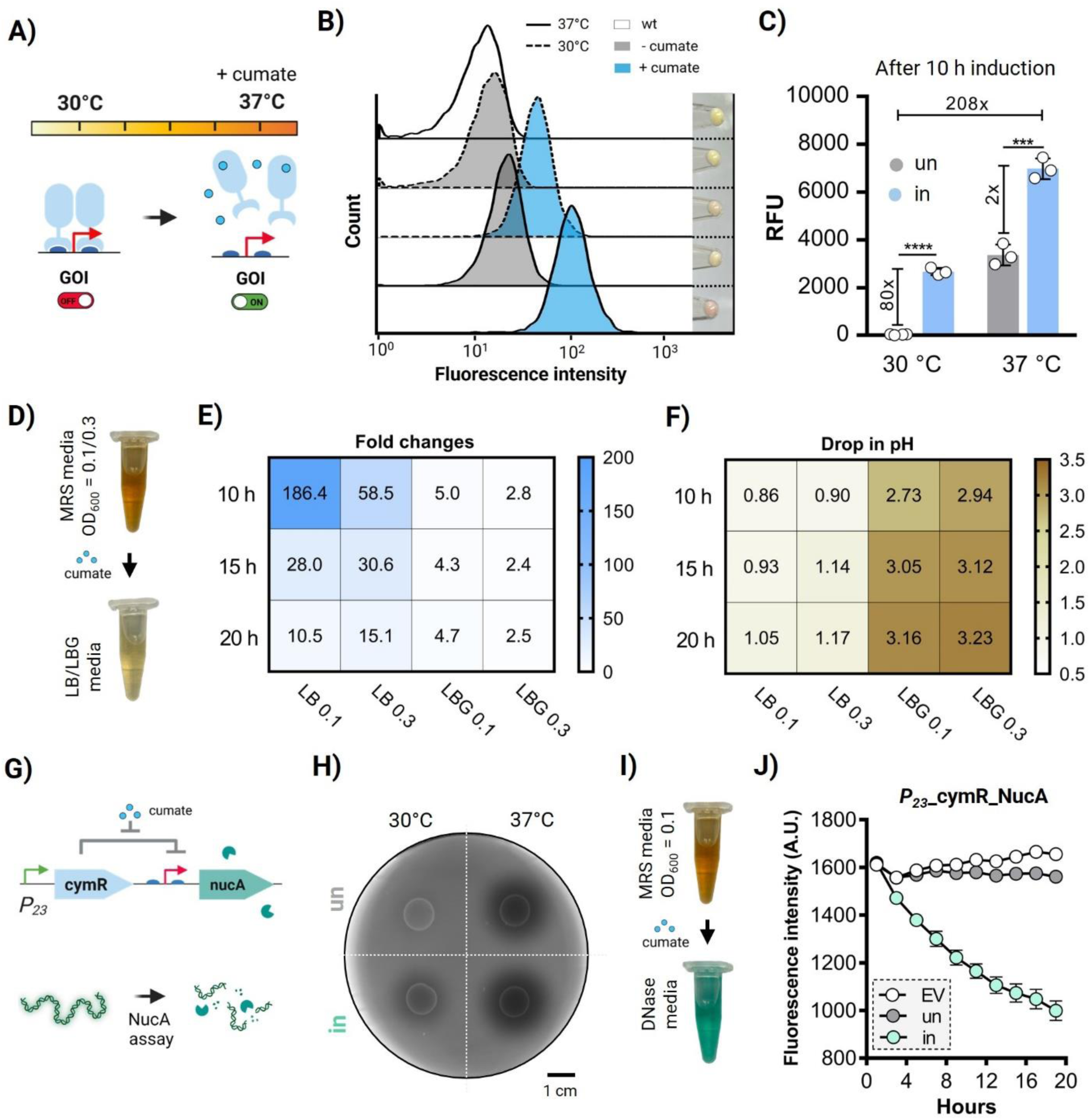
**A)** Scheme depicting the thermally aided switching strategy involving both temperature and cumate. Proper turning on of the gene is achieved via both the incubation at 37°C and the addition of cumate. **B)** Histograms showing the fluorescence intensity (after 8 hours) of the *P_23_*_cymR_mCherry variant at 30 and 37°and with and without the addition of cumate. Bacterial pellets are shown on the right of each condition. **C)** RFU values corresponding to hour 10 after induction for the *P_23_*_cymR_mCherry variant when induced at 37°C and 30°C (n = 3, mean ± SD). “un” = uninduced condition, “in” = induced condition (5 µM of cumate). **D)** Illustration showing the procedure of bacterial growth and subculturing. Bacteria were subcultured in MRS medium and grown until an OD_600nm_ of either 0.1 or 0.3 and then transferred to LB or LBG media with and without cumate. **E)** Heatmap plot depicting the fold changes between the induced and uninduced conditions in LB and LBG media with an initial OD_600nm_ of 0.1 and 0.3 after 10, 15, and 20 hours of incubation at 37°C. Bacteria were induced with 5 µM of cumate (n = 4, mean ± SD). **F)** Heatmap depicting the drop in pH after 10, 15, and 20 hours of growth in either LB or LBG media at initial OD_600nm_ of 0.1 and 0.3 and at 37°C. The pH values were deducted from the initial pH value (0 h) of each condition (medium and OD600). **G**) Scheme depicting the cumate genetic switch controlling NucA expression, which can be quantified via the NucA assay. **H**) 10 µL of *P_23_*_cymR_NucA bacteria, uninduced and induced for 8 hours in MRS medium at both 30 and 37°C, were spotted in a DNase agar plate and incubated at 37°C for 40 hours. “un” = uninduced condition, “in” = induced condition (5 µM of cumate). **I)** Illustration showing the procedure of bacterial growth and subculturing. Bacteria were subcultured in MRS medium and grown at an OD_600nm_ of either 0.1 and then transferred to DNase medium with and without cumate. **J)** Plot showing the drop in the fluorescence intensity as a result of the NucA secretion (n = 4, mean ± SD). EV = empty vector, un” = uninduced condition, “in” = induced condition (5 µM of cumate).

We first evaluated the influence of temperature on the *P_23_*_cymR_mCherry variant through flow cytometry to confirm that the responses to temperature and cumate occurred across the entire population. Cultures prepared at a bacterial density of 0.01 OD_600nm_ were induced with 5 µM of cumate and incubated at both 30 and 37°C for 8 hours. Results showed low leakiness of the switch at 30°C without induction and prominent levels of mCherry expression at 37°C (**Figure 2B**). Besides, induction at 30°C also worked very efficiently, with the entire population switching to higher levels of fluorescence intensity. Thermally aided switching was even visible in the coloration of the bacterial pellets (**Figure 2B**), which increased with both higher temperature and cumate induction.

We then estimated the kinetic performance of the genetic switch at both temperatures with and without cumate. The *P_23_*_cymR_mCherry variant exhibited a considerable delay in the start of leaky expression when grown at 30°C (>11 hours) instead of 37°C (>8 hours) while maintaining notable induced expression in the presence of cumate by these time points (**Figure 2C**). These time points corresponded to a higher bacterial density towards the end of the log phase at 30°C (∼0.7 OD_600nm_) compared to 37°C (∼0.4 OD_600nm_) (**Figure S5B** and **S5C**). This suggested that at 30°C, leakiness is suppressed until a slightly higher biomass compared to 37°C, although induced protein production at 30°C is lower.

Accordingly, after 10 hours of induction, we calculated an 80-fold difference between uninduced and induced states at 30°C, and a 208-fold difference between the uninduced state at 30°C and the induced state at 37°C (**Figure 2C**).

Alternatively, we tested the cumate switch in a non-optimal medium to limit the growth of the bacteria while maintaining its protein production capability. Bacteria were grown until an OD_600nm_ of either 0.1 or 0.3 in MRS medium and then transferred into either LB medium or LB medium supplemented with 0.1 M of glucose (LBG) (**Figure 2D**). Minimal leakiness was detected in LB medium after 15 hours for both 0.1 and 0.3 initial OD_600nm_ whereas strong leaky expression started after 6 and 4 hours for the 0.1 OD_600nm_ and 0.3 OD_600nm_ bacteria grown in LBG medium, respectively (**Figure S6A**). When glucose was added, bacterial growth was significantly higher (**Figure S6B**), confirming that the leakiness of the cumate switch is linked to the exponential growth of the bacteria. Since lactobacillus growth and metabolism are typically associated with acid formation, we suspected that pH may play a role in disrupting the activity of the repressor. Accordingly, we measured changes in pH in both LB and LBG media after 20 h of bacterial growth. As suspected, while both media had an initial pH of nearly 6.8, this value dropped much more in LBG (3.5 – 3.6) than LB (5.6-5.7) medium (**Figure S6C** and **S6D**). Consequently, we observed an inverse correlation between the fold-change of the cumate switch and the drop in pH mediated by bacterial growth (**Figures 2E**, **2F, S6C,** and **S6D**). The same inverse correlation between a pH drop and fold-change was observed when the bacteria were grown in the optimal MRS medium, at 30 and 37 °C (**Figure S7A, S7B,** and **S7C**). These results suggested a strong correlation between bacterial growth, pH drop, and leakiness of the cumate switch. We speculate that a drop in pH could alter the conformation of the repressor protein and/or its binding affinity with the operator sequence [39–41].This may be a key factor affecting the performance of other poorly performing genetic switches in lactobacilli that have been previously reported in the literature [16, 17]. It should be noted that *L. plantarum* has been reported to maintain a higher cytoplasmic pH compared to surrounding acidic media by 0.5 – 1 pH units using proton pumps [42]. Thus, the pH we measured in the medium can only be considered as an indication of a pH change in the cytoplasm but cannot precisely indicate cytoplasmic pH values. Nevertheless, our results indicated that the performance of the cumate switch can be improved by growing bacteria in an environment that limits growth to prevent a drop in pH.

Since the cumate switch functions best under growth-limited conditions, intracellular protein production capacity would be limited by the amount of biomass in a culture. To overcome this limitation, we tested the possibility of using the switch to mediate protein secretion, which would enable more protein to accumulate in the culture from a given biomass. Protein secretion is also of great interest for industrial and medical applications since it allows for easier protein purification and programming bacteria to exert extracellular bioactivity [43–45]. For this, we used NucA, a nuclease from *Staphylococcus aureus*, as a reporter protein for secretion since there is a simple assay to measure its activity [46–48]. The assay is based on the degradation of fluorescently labeled DNA by NucA. Using this method, we previously reported the Lp_3050 signal peptide to be highly efficient for secreting NucA with the *P_tec_* promoter [12]. Accordingly, we cloned the Lp_3050_NucA gene under the control of the cumate switch, generating the *P_23_*_cymR_NucA variant (**Figure 2G**).

Based on the results with the mCherry expression, we expected that at 30°C without cumate, there would be low levels of NucA secretion whereas, at 37 °C with cumate, secretion of NucA would be higher. We evaluated thermally aided switching in both DNase agar and liquid medium, which contained fluorescently labeled DNA. In the agar experiment, 10 µL of induced and uninduced bacteria grown in MRS until an OD_600nm_ of 1 at both 30°C and 37°C were spotted on DNase agar and incubated at 37°C for roughly 40 hours. We could not discern a halo around the spotted area of bacteria grown at 30°C without cumate. On the other hand, there were clearly visible dark halos for the other three conditions, with the darkest and widest ones for the bacteria grown at 37°C with and without cumate (**Figure 2H** and **S8A**). These results indicate that at 37°C, the cumate switch is also leaky for NucA, which matches the results obtained with mCherry. Image analysis of the fluorescence intensity drop in the halo revealed that, at 37°C, the halo was slightly darker with cumate than the one without (**Figure S8B**). For better quantification, the bacteria were grown and induced in the same way but then transferred into liquid DNase medium with or without cumate and incubated at either 30°C or 37°C. After 24 hours, the fluorescence of the medium was measured, and the drop in fluorescence was correlated with a standard curve to estimate the concentration of secreted NucA. At 30°C without cumate, NucA levels were below the detection limit, while over 1 nM of cumate was secreted from the bacteria with cumate. At 37°C, roughly 9 and 11 nM were detected in the absence and presence of cumate. respectively (**Figure S8C**). In contrast, the performance of the cumate switch greatly improves under low bacterial density conditions. When the bacteria were grown to an OD_600nm_ of 0.1 in MRS medium and then transferred to DNase medium (**Figure 2I)**, secretion of NucA under uninduced conditions was negligible for 20 h (**Figure 2J**). On the other hand, a prominent and continuous drop in fluorescence was detected upon the addition of cumate over 20 h, suggesting steady production and secretion of NucA (**Figure 2J**).

Overall, the results with both mCherry and NucA proved that thermally aided switching and the use of nutrient-limiting medium can be efficient approaches to tightly regulate gene expression under the cumate switch in *L. plantarum* when grown in culture.

### High-performance encapsulated genetic switches

In our previously reported alginate core-shell encapsulation format for genetically engineered *L. plantarum* (PEARL) [12], we observed steady protein production and secretion for 14 days without outgrowth of the bacteria. The results from that study suggested that the encapsulated bacteria stopped growing after a day or two but remained metabolically active to produce and secrete *P_tec_*-driven recombinant proteins at a rate corresponding to log-phase in liquid cultures. From the results of the current study suggesting the best performance of the cumate switch in the early to mid-log phase, we speculated that encapsulation in the PEARL format could prolong this performance. Since cumate is a small-molecule inducer, it was expected to diffuse inside the PEARLs and activate protein production in the encapsulated bacteria **(Figure 3A**). After fabricating PEARLs with the *P_23_*_cymR_mCherry variant, cumate was added to the medium surrounding them and fluorescence was measured over 3 days. The expression of mCherry in these induced PEARLs was detected after 4 hours, slowly increased to ∼12 hours then rapidly increased to nearly 24 hours (**Figure 3B**). The initial slow increase in expression may be due to the bacterial metabolism simultaneously supporting growth and mCherry production during this time. This is supported by our previously reported observation that *L. plantarum* growth in PEARLs plateaus around 12 hours [12]. Beyond this point, the metabolic resources in the bacteria are forced predominantly towards mCherry production. The slight drop in fluorescence after 24 hours suggests a saturation of the expressed protein within the bacteria and possibly bacterial death due to it (**Figure 3B**). More striking was the observation that leaky mCherry expression in the uninduced PEARLs was only observed after 15 hours, increased a little to ∼20 hours, and plateaued beyond this time point. On calculating the fold change between the induced and uninduced PEARLs, the highest value was surprisingly observed at 10.5 hours (92.5x), possibly due to negligible leaky expression, and after 24 hours, it stabilized between 25-35x (**Figure S9A**). These results confirmed our hypothesis that encapsulation can help to reduce the leakiness of the cumate switch.

**Figure 3:**
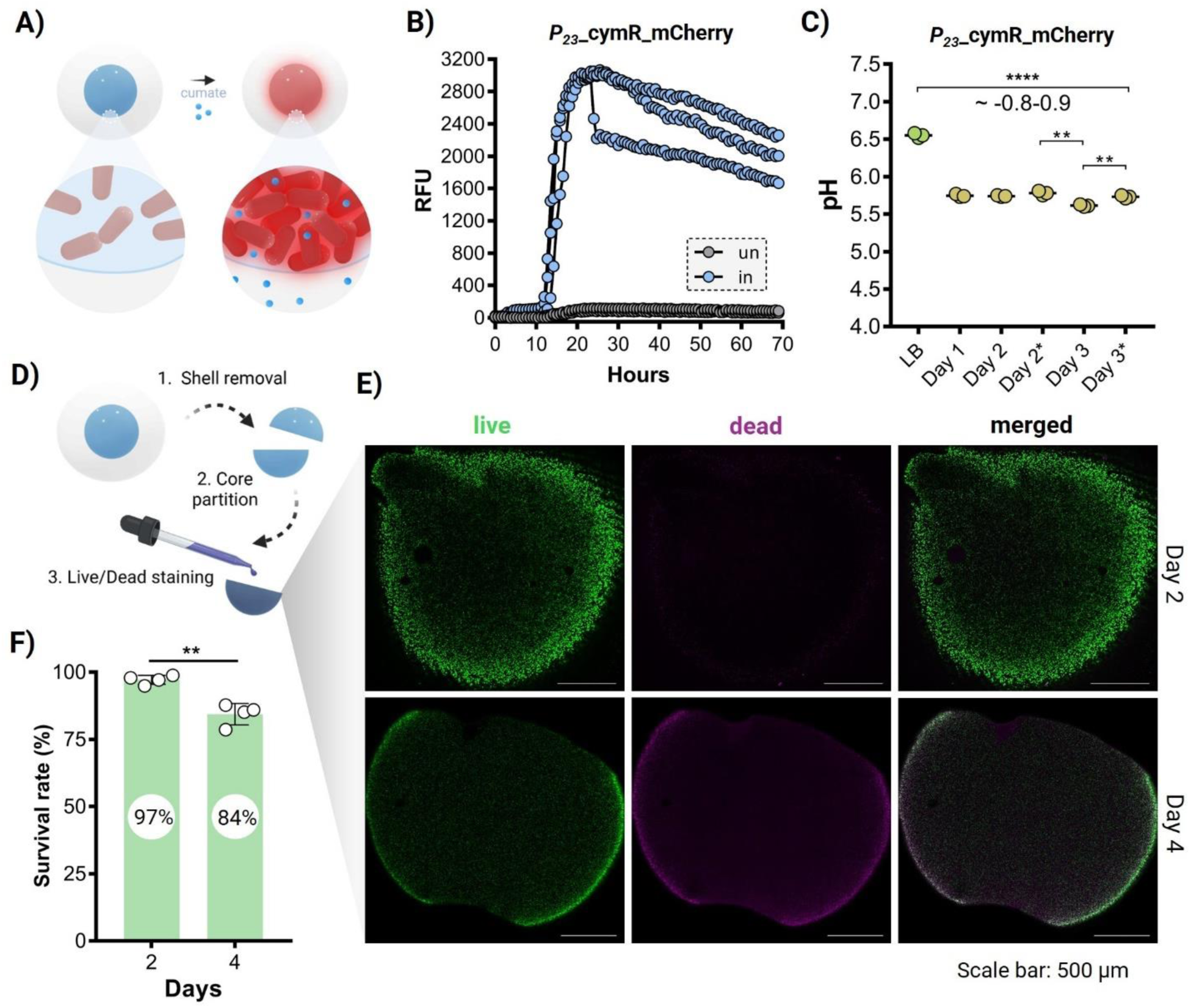
**A**) Scheme showing the induction of *L. plantarum* bacteria harboring the cumate switch when encapsulated in PEARLs. Upon induction, the core of the PEARLs turns red due to bacterial activation and subsequent mCherry intracellular production. **B)** RFU for the *P_23_*_cymR_mCherry variant when encapsulated in PEARLs and incubated at 37°C for 24 hours (n = 3, mean ± SD). “un” = uninduced condition, “in” = induced condition (100 µM of cumate). **C**) Plot showing the significant pH drop in the LB medium when PEARLs were incubated for 1 to 3 days at 37 °C. * = fresh LB medium was added every 24 hours. **D**) Illustration showing how PEARLs were processed before the live/dead imaging. The shell was carefully detached from the core, which was portioned in half prior to being subjected to the live/dead staining for 30 minutes at room temperature in static. The stained cores were washed with PBS before imaging. **E**) Representative images of the live/dead staining of the cores of PEARLs after 2 and 4 days of incubation. **F**) Plot showing the percentage of bacterial survival in the PEARLs after 2 and 4 days (n = 4, mean ± SD).

One possible explanation would be due to the stable maintenance of the pH. We measured the pH in the medium surrounding the PEARLs over three consecutive days of incubation at 37 °C, and the drop was 0.80, 0.81, and 0.93 after 1, 2, and 3 days of incubation, respectively (**Figure 3C**). Similar drops, 0.77 and 0.82, were measured when the media were refreshed each day after the first 24 hours (**Figure 3C**). These magnitudes of pH drops are similar to those seen with non-encapsulated cells in LB medium, where the highest fold changes were recorded (**Figure 2E**). During this time, the viability of the bacteria, determined by live/dead staining and confocal microscopy (**Figure 3D** and **3E**), revealed a slight drop from an average of 97% viable on day 2 to 84% on day 4 (**Figure 3F**). This could be contributing to the drop in fluorescence seen after day 1 (**Figure 3B**).

When comparing the behavior *P_23_*_cymR_mCherry with *P_tec_*_cymR_P_mCherry in the PEARL format, we found that both performed very similarly with *P_tec_*_cymR_P_mCherry being a bit leakier (**Figure S9B** and **S9C**). In comparison, the pSIP_409 variant in the PEARL format exhibited expectedly negligible levels of uninduced leaky expression and low levels of induced mCherry production - ∼37 and ∼50 times lower than the *P_23_*_cymR_mCherry and the *P_tec_*_cymR_P_mCherry variants, respectively (**Figure S9D**). Thus, encapsulation enhances the performance of the cumate switch by suppressing leaky expression but does not improve the performance of the pSIP_409 switch, since it has no influence over the promoter strength.

As an additional demonstration, we tested the performance of the cumate switch in a mammalian cell culture medium, DMEM. This is a sub-optimal medium for *L. plantarum* growth and we have previously shown the possibility of secreting therapeutic proteins in it using PEARLs [12]. As anticipated, the cumate switch also performed really well in this medium with strong inducible expression at 24 h and poor uninduced expression (**Figure S9E**). This indicates the possibility of using switchable PEARLs to regulate the production of proteins under conditions suitable for mammalian cells.

### Switchable protein secretion from PEARLs

Since the PEARL format supported stable switching of mCherry production, we proceeded with the encapsulation of the *P_23_*_cymR_NucA variant to test switchable protein secretion. For this, we incubated the PEARLs in liquid DNase medium and measured changes in its fluorescence intensity over time (**Figure 4A**). Over 9 days, negligible leaky expression was observed in uninduced conditions, whereas a prominent fluorescence drop was detected on all days, indicating steady secretion and release of NucA from the PEARLs (**Figure 4B**). The magnitude of the fluorescence intensity drop reduced from ∼60% to ∼40% over the first five days, after which it remained steady till day 9. This may be due to a loss in bacterial viability over the first five days and then stabilization of the encapsulated population. Next, we tested four different switching scenarios over 2 days in the absence and presence of cumate: i) OFF-OFF - PEARLs incubated in DNase medium without cumate for 1 day, then washed with sterile Milli-Q water, followed by another entire day incubation with fresh DNase medium without cumate. ii) OFF-ON - day 1 without cumate and day 2 with cumate. iii) ON-OFF - day 1 with cumate and day 2 without cumate. iv) ON-ON - day 1 with cumate and day 2 with cumate. The OFF-OFF scenario showed that the genetic switch remained in the OFF state on both days with no detectable NucA secretion (same as control), which means that encapsulation effectively prevented leaky expression (**Figure 4C**, top left). The OFF-ON scenario also worked well, with encapsulated bacteria being activated only on the second day upon the addition of cumate in the fresh DNase medium (**Figure 4C**, bottom left). This finding is particularly interesting from the ELM perspective since it highlights that encapsulated bacteria could remain inactive for hours and then be activated on demand. For the ON-ON scenario, as expected, we detected a strong drop in fluorescence for the first and second day as well, confirming that the presence of cumate in the medium sustained NucA secretion from the bacteria even on the second day (**Figure 4C**, top right). Finally, in the ON-OFF scenario, while there was strong secretion on the first day, switching did not effectively occur on the second day. There was a slight decrease in the magnitude to which fluorescence in the DNase medium dropped (53% and 38% drop in fluorescence compared to the control on the first and second day, correspondingly), but the inactivation was not as sharp as the activation in the OFF-ON scenario (**Figure 4C**, bottom right). This could be due to retention of cumate within the PEARLs, keeping the switch active, slow release of accumulated NucA from the alginate, or continued secretion of NucA that was already produced in the bacteria. Most likely, all three phenomena influenced this outcome. Nevertheless, the result indicates that encapsulation can prolong the ON state beyond the period when the inducer is present. This is desirable in scenarios like living therapeutics for prolonged drug release with a short activation signal.

**Figure 4:**
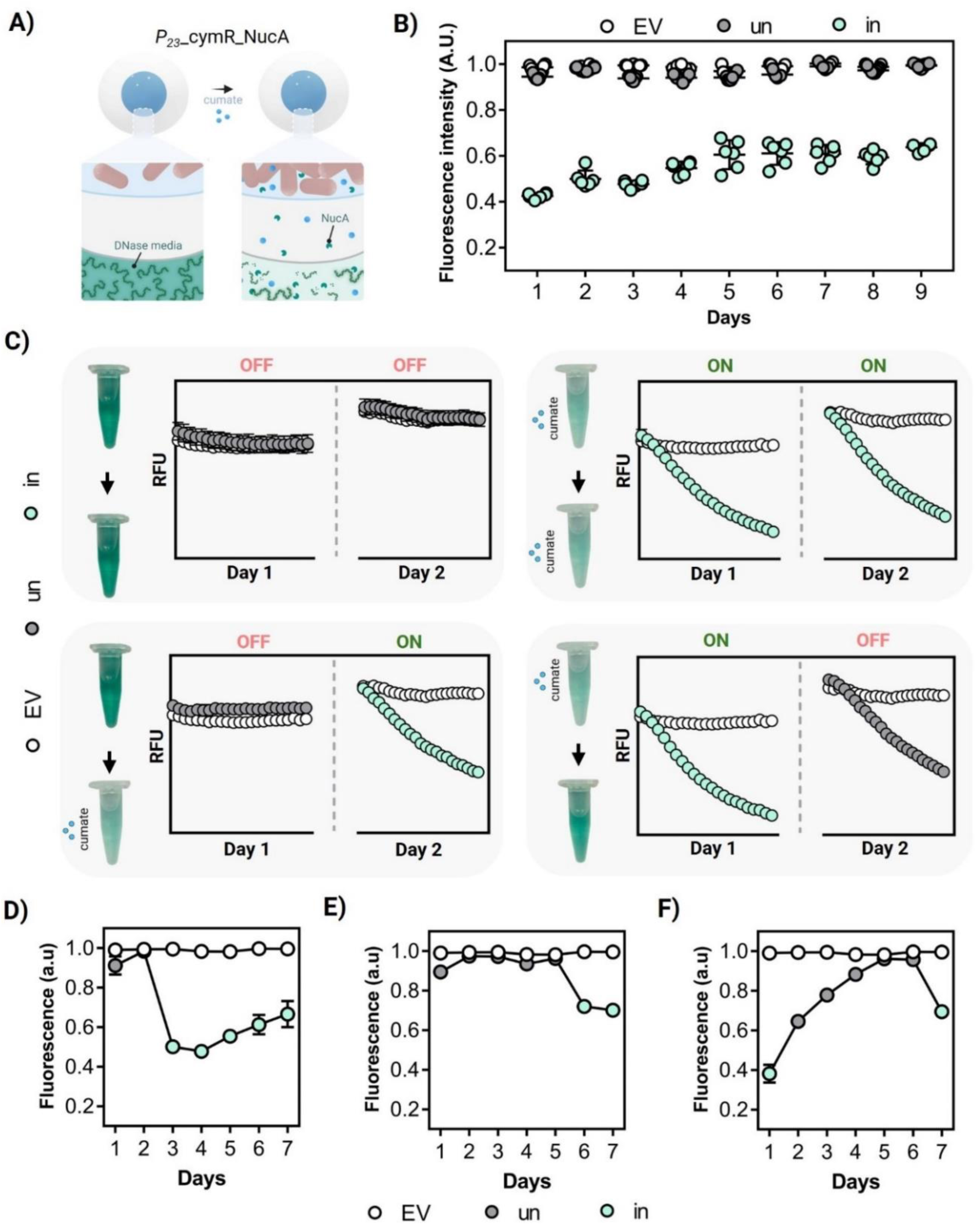
**A)** Scheme showing the NucA production regulated by the cumate switch when bacteria are encapsulated in PEARLs. The cumate diffuses inside the core and activates the bacteria, which triggers the secretion of NucA into the shell. The diffusion of NucA into the DNase medium results in the discoloration of the medium due to NucA activity. **B**) Plot showing the detection of secreted NucA (as a drop in fluorescence intensity) from the supernatant of PEARLs from day 1 to day 9 (n = 6, mean ± SD). EV = empty vector, “un” = uninduced condition, “in” = induced condition (100 µM of cumate). **C**) *P_23_*_cymR_NucA ON and OFF switches depending on the incubation of the PEARLs in DNase medium with or without cumate on day 1 and day 2. PEARLs were washed between days 1 and 2 and measurements are shown as RFUs (n = 4, mean ± SD). **D**) Extended OFF-ON switch where PEARLs were incubated for two days in cumate-free DNase medium before induction with cumate for 5 consecutive days. The PEARLs were washed every day before adding fresh medium (n = 4, mean ± SD). The images of the tubes with DNase medium are only displayed for better clarity of the switches. **E**) Extended OFF-ON switch where PEARLs were incubated for five days in cumate-free DNase medium before induction with cumate for 2 consecutive days. The PEARLs were washed every day before adding fresh medium (n = 4, mean ± SD). **F**) Extended ON-OFF switch where PEARLs were incubated with DNase medium with cumate for 1 day, followed by 5 consecutive days in cumate-free DNase medium and a final incubation day in DNase medium with cumate. The PEARLs were washed every day before adding fresh medium (n = 4, mean ± SD). EV = empty vector, “un” = uninduced condition, “in” = induced condition (100 µM of cumate).

Based on these results, we studied the OFF-ON and the ON-OFF switching conditions in more detail over seven days. It was possible to activate NucA secretion from encapsulated bacteria using cumate after leaving them in the OFF state for 2 and 5 days (**Figures 4D** and **4E**). Notably, the level of NucA release was lower on the later days, with fluorescence intensity drops of the DNase medium reaching 30-40% on day 7. As for the ON-OFF switching condition, we observed that PEARLs activated on day 0 and turned OFF on day 1 kept releasing NucA for up to four days. Importantly, by day 5, almost no NucA was observed in the medium, indicating that the cumate switch retained its tight control over expression even after being activated once. Moreover, the switch could be activated once more on day 6 resulting in a ∼30% drop in DNase medium fluorescence intensity by day 7 (**Figure 4F**). These results indicated the possibility of switching between different states in PEARLs, which is highly desirable for developing smart ELMs [28].

### Dual switchable PEARLs

Altogether, bacterial encapsulation within PEARLs enhanced the performance of the cumate switch by reducing the non-induced state leakiness while maintaining activity in the induced state. Therefore, we tested whether it would work for the highly leaky vanillate switch (**Figure 1C** and **S1C**). Accordingly, we observed a considerably enhanced performance of the vanillate switch when bacteria were encapsulated within PEARLs, with fold changes close to 15x after 20 hours of induction (**Figure 5A** and **S10A**). These results were validated by fluorescence imaging, where red fluorescence was observed only in the cores of the induced PEARLs (**Figure 5B**). On the other hand, kinetic analysis of mCherry production in non-encapsulated bacteria with the vanillate switch yielded very poor fold changes that were consistently below 1.5x (**Figure S10B**). Thus, alginate bead encapsulation enhanced the performance of the vanillate switch by an order of magnitude.

**Figure 5:**
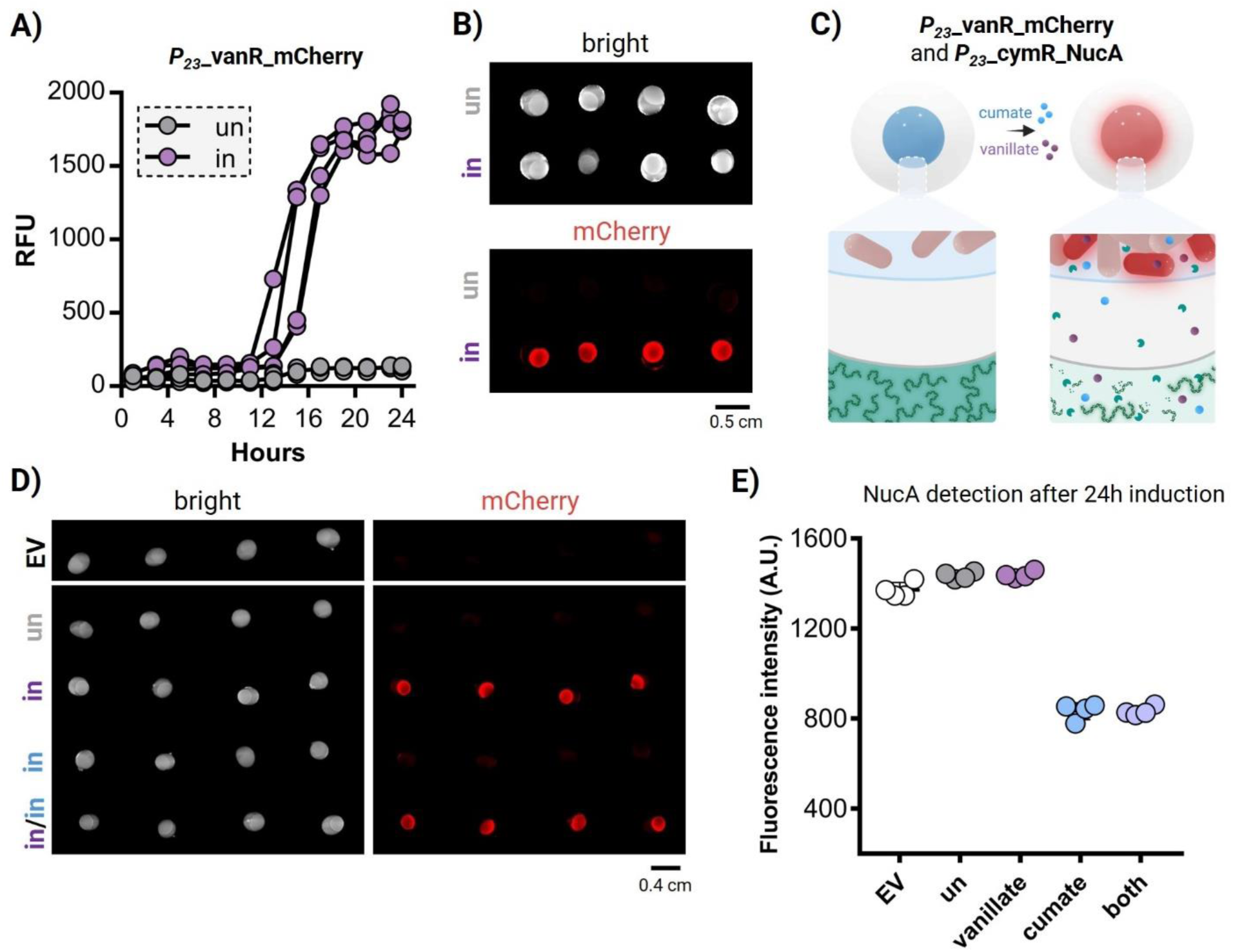
**A)** RFU values for the induced and uninduced states of the *P_23_*_vanR_mCherry variant when encapsulated in PEARLs and grown at 37°C for 23 hours (n = 4, mean ± SD). “un” = uninduced condition, “in” = induced condition (100 µM of vanillate). **B**) BioRad Gel Documentation System images of the induced and uninduced *P_23_*_vanR_mCherry-based PEARLs after 23 hours of growth at 37°C (n = 4, mean ± SD). “un” = uninduced condition, “in” = induced condition (100 µM of vanillate). **C**) Scheme depicting the co-encapsulation of bacteria producing mCherry via the vanillate switch and secreting NucA via the cumate switch. Both inductions can be detected from the PEARLs by assessing the red fluorescence increase in the core of the PEARLs and the drop in the fluorescence in the DNase medium surrounding the PEARLs. **D**) Red fluorescence captured from the PEARLs (*P_23_*_cymR_NucA and *P_23_*_vanR_mCherry) that were incubated without any inducer, only with vanillate, only with cumate and with both inducers at 37°C for 24 hours (n = 4, mean ± SD). Images were taken with the BioRad Gel Documentation System. EV = empty vector, “un” = uninduced condition, “in” = induced condition (100 µM of cumate, 100 µM of vanillate, or both). **E**) Drop in fluorescence triggered by NucA secretion into the DNase medium from the PEARLs. Empty Vector PEARLs were kept as negative control (n = 4, mean ± SD).

Based on these promising results, we attempted co-encapsulation of the two *L. plantarum* strains engineered with cumate (*P_23_*_cymR_NucA) and vanillate (*P_23_*_vanR_mCherry) switches **(Figure 5C**). Encapsulation of bacteria in hydrogels offers the unique opportunity to establish stable co-cultures and thereby create multi-functional ELMs [49–51]. The dual switchable PEARLs were induced individually with vanillate and cumate and with both inducers simultaneously. Fluorescence imaging revealed that only the PEARLs that were induced with vanillate showed detectable levels of mCherry expression (**Figure 5D**). In parallel, their NucA secretion was assessed by monitoring the drop in fluorescence in DNase medium, which was observed only in the PEARLs induced with cumate (**Figure 5E**). Furthermore, both populations can also be induced simultaneously when both inducers are added to the medium (**Figure 5D** and **5E**). This demonstrated that multi-switching and multi-functional PEARLs can be created by encapsulating different engineered *L. plantarum* strains.

### Cumate genetic switch in micro-PEARLs

Encouraged by these results, we proceeded to develop the PEARL system into a more compact format better suited for different applications. Furthermore, the PEARL format faces other limitations since it is a time-consuming manual approach that can lead to fabrication errors and has poor scalability. Therefore, we adopted a more automated, time-saving, up-scalable approach to produce a higher number of beads with smaller sizes [52, 53]. For this, we used an electrospray microencapsulator B-390/B-395 (Buchi) to generate core-shell alginate micro-beads (micro-PEARLs) that ranged in sizes from 100 µm to 500 µm. In these micro-PEARLs, we encapsulated a single *L. plantarum* strain that constitutively produced a red fluorescent protein, mScarlet3, and secreted NucA (**Figure 6A**). Hence, we could simultaneously monitor gene expression within the micro-PEARLs by tracking red fluorescence production and assessing protein secretion through the DNase assay. After incubating the micro-PEARLs for 24 hours on a DNase agar plate, we detected a prominent dark halo around the set of micro-PEARLs, and red fluorescence in their core. Furthermore, the halo increased in size after 5 days, suggesting continued secretion of NucA over time (**Figure 6B, S11A**). Moreover, bacterial growth within the cores could be observed even by eye after incubating the micro-PEARLs in 2-mL tubes with MRS medium for 24 hours (**Figure 6C**).

**Figure 6:**
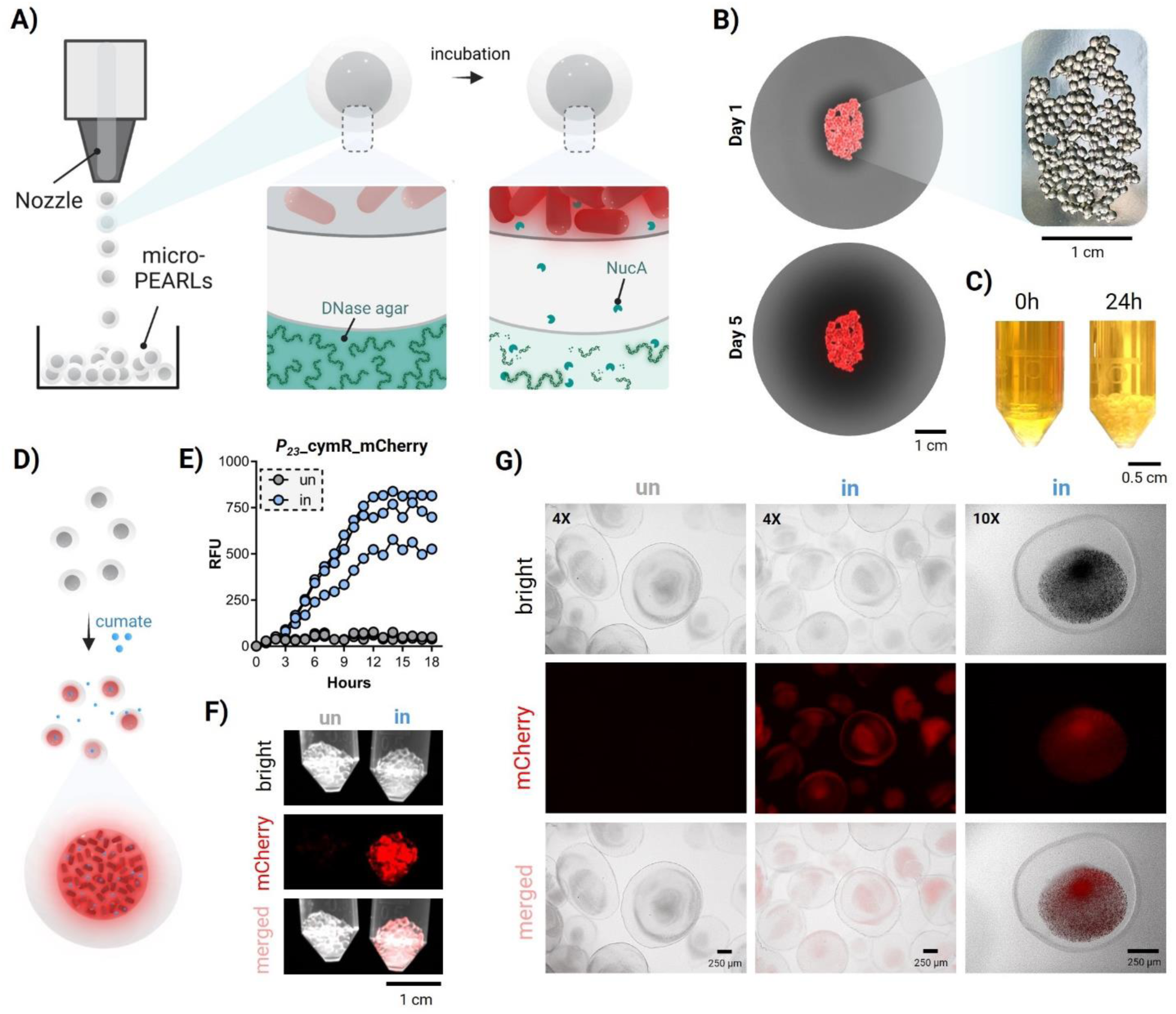
**A)** Scheme showing micro-PEARL fabrication with the microencapsulator. The micro-PEARLs contain bacteria that produce mScarlet3 and secrete NucA. After incubation, the core of the micro-PEARLs turns red, and the fluorescence of the DNase agar fades away due to the degradation of fluorescently labeled DNA. **B)** DNase agar plates with micro-PEARLs after 1 and 5 days of incubation at 37°C. Images taken using the BioRad Gel Documentation System. **C)** 2 mL tubes with micro-PEARLs incubated in MRS medium after 0 and 24 hours. The diameter of the 2- mL tube is 11 mm. **D)** Scheme of the cumate induction in micro-PEARLs. Upon induction, the core of the micro-PEARLs turns red due to mCherry intracellular production. **E**) RFU values of the *P_23_*_cymR_mCherry variant when encapsulated in micro-PEARLs and induced with cumate for 19 hours (n = 3, mean ± SD). “un” = uninduced condition, “in” = induced condition (100 µM of cumate). **F**) BioRad Gel Documentation System images of uninduced (un) and induced (in) micro-PEALSs after 24 hours of incubation at 37 °C. Induction was done with 100 µM of cumate. **G**) Microscopy images, at both 10X and 4X, of uninduced (un) and induced (in) micro-PEARLs after 24 hours of incubation at 37 °C. Induction was done with 100 µM of cumate.

Next, we evaluated if these micro-PEARLs could be induced upon the addition of cumate in the medium. For that, we used the *P_23_*_cymR_mcherry variant, which would ideally produce mCherry in the core of the micro-PEARLs when cumate is added to the medium (**Figure 6D**). As expected, it was possible to track the induction over time with a clear increase in red fluorescence only in the presence of cumate (**Figure 6E**). Kinetic analysis over 24 hours clearly revealed that even in the micro-PEARLs format, encapsulation ensured almost completely uninduced repression of mCherry expression. This difference was even visible through macroscopic fluorescence imaging (**Figure 6F**). Microscopic analysis of the micro-PEARLs revealed mCherry fluorescence predominantly in the cores and only in those that were induced by cumate (**Figure 6G**, **S11B,** and **S11C**). These results validate that the more automated, time-saving, effective, and compact encapsulation method for *L. plantarum* allows the bacteria to grow efficiently while maintaining them in an active and cumate-switchable state.

## CONCLUSIONS

In this study, we have successfully modified *L. plantarum*’s strongest constitutive promoter, *P_tec_*, into a genetic switch responsive to cumate. However, the switch suffered from leakiness of gene expression associated with bacterial growth and a drop in pH, especially when the bacteria entered the late log phase. Due to the natural capacity of *P_tec_* to be more active at higher temperatures, we explored a workaround to reduce the leakiness of the cumate switch through thermally aided switching, in which the uninduced state was maintained at 30°C and cumate induction was performed at 37°C. The performance of the cumate switch was also increased by subculturing the bacteria in non-optimal medium that limited growth but supported protein production. However, a more promising strategy to enhance the performance of the switch was discovered through the encapsulation of the engineered bacteria in alginate beads to form PEARLs. This surprisingly suppressed leaky expression of the cumate switch while supporting high levels of protein production and secretion. This effect extended beyond the cumate switch to the much leakier vanillate switch and even sustained the performance of both switches in a co-culture setting. Furthermore, the encapsulation-enhanced switching was not dependent on the size of the encapsulation matrix as the effect persisted even in micro-PEARLs, which can be rapidly produced in an automated manner. We hypothesize that encapsulation within alginate mechanically and metabolically restricts bacterial growth, which maintains the pH of the environment more stably and favors the repression of the leaky expression.

These encapsulation-enhanced switches are promising for the development of responsive ELMs for medical, environmental, and industrial applications [54–56]. By combining this effect with mechanically switchable smart materials, it may be possible to control the performance of such genetic switches by exerting *in situ* changes in the mechanical properties of the material.

## EXPERIMENTAL SECTION

### Strains, media and plasmid vector

*L. plantarum* WCFS1 was used in this study. The strain was grown in De Man, Rogosa, and Sharpe (MRS) medium (Carl Roth GmbH, Germany, art. no. X924.1). The strain was incubated at either 30° or 37°C (indicated in each case), and 250 revolutions per minute (rpm). The medium of the engineered strains was supplemented with 10 μL/mL of erythromycin (Carl Roth GmbH, Germany, art. No. 4166.2). For the cloning of all the plasmids, NEB DH5-α competent *E. coli* cells were used (New England Biolabs GmbH, Germany, art. No. C2987). Engineered *E. coli* was grown in Luria-Bertani (LB) medium (Carl Roth GmbH, art. No. X968.1) supplemented with 200 μg/mL of erythromycin at 37°C and 250 rpm shaking conditions. The pTec_mCherry plasmid (Addgene plasmid #229145) was the backbone vector for all the plasmids used in this work. The backbone of this plasmid is derived from the pLp_3050sNuc plasmid (Addgene plasmid # 122030).

### Molecular cloning reagents

Primers were synthesized by Eurofins Genomics GmbH (Köln, Germany). All the primers used in this study are listed in **Table S1**. Synthetic gene fragments were ordered as eBlocks from Integrated DNA Technologies (IDT) (Coralville, USA). The codon optimization tool of IDT was used to codon-optimize the eBlocks (selecting the option for *Lactobacillus acidophilus*). All the eBlocks used in this study are listed in **Figure S0**. For the polymerase chain reaction (PCR), Q5 High Fidelity 2X Master Mix (NEB GmbH, Germany, No. M0492S) was used. 1 kb Plus DNA Ladder (Catalog Number 10787018) was purchased from ThermoFisher Scientific (Germany) and used as a reference in gel electrophoresis. The Wizard® SV Gel and PCR Clean-Up System (Promega GmbH, Germany, Art. No. A9282) was used to purify the PCR products. DNA assembly was performed using the HiFi Assembly Master Mix (NEB GmbH, Germany, Art. No. E5520S). The Qiagen GmbH (Hilden, Germany) plasmid extraction kit was used to lyse and extract the plasmids from *E. coli* DH5α.

### Plasmid construction

The eBlocks were designed with compatible ends (20 base-pair overhangs) to the backbone vector. Each eBlock was resuspended with 20 µL of previously autoclaved Milli-Q water and used directly in the HiFi assembly reaction following the manufacturer’s protocol. Right after, the assembled products were transformed into NEB DH5-α *E. coli* competent cells following the manufacturer’s heat shock protocol except for the NucA cloning, where bacteria were grown at 28°C instead of 37°C. Plasmids were extracted from *E. coli* DH5-α cells and used for transforming *L. plantarum* WCFS1 after sequence verification by Sanger sequencing (Eurofins Genomics GmbH (Köln, Germany)).

### *L. plantarum* WCFS1 competent cell preparation and DNA transformation

The preparation of *L. plantarum* WCSF1 electrocompetent cells and the plasmid DNA transformation were done as described in [57]. In brief, *L. plantarum* WCFS1 was inoculated in 5 mL of MRS medium and cultured overnight at 37°C and 250 rpm. After overnight growth, 1 mL of the culture was mixed with 20 mL of MRS medium and 5 mL of 1% (w/v) glycine and incubated at 37°C and 250 rpm for 4 hours approximately until OD_600nm_ reached roughly 1. Next, the cells were pelleted down by centrifuging at 4000 rpm (3363 g) for 10 min at 4°C, followed by two washes with 5 mL of ice-cold 10 mM MgCl2 and two additional washes with 5 mL of ice-cold Sac/Gly solution [1M sucrose mixed with 10% (v/v) glycerol in a 1:1 (v/v) ratio]. After that, cells were resuspended in 500 μL of Sac/Gly solution, and 60 μL aliquots of competent cells were prepared for DNA transformation. For the electroporation, 1 μg of plasmid DNA was added to a competent cells aliquot prior to transferring the mixture into an ice-cold 2 mm gap electroporation cuvette (Bio-Rad Laboratories GmbH, Germany). After 10 minutes of incubation on ice, a single pulse at 1.8 kV was applied, followed by the addition of 1 mL of MRS medium. The mixture was incubated at 37°C and 250 rpm for 3 hours. Finally, all the cells were plated on MRS Agar supplemented with 10 μg/mL of Erythromycin and incubated at 37°C for 48 hours.

### Gene expression switches induction

Vanillic acid (vanillate) and 4-isopropylbenzoicacid (cumate) were purchased at Sigma Aldrich (Catalog number H36001 and 268402, respectively). Anhydrotetracycline (aTc) was purchased at Cayman Chemical (Item no. 10009542). All inducers were resuspended in pure ethanol. The concentrations used for inductions were: 50, 100, or 200 µM of vanillate for the vanillate switch, 0.2, 0.5, 1, or 4 µM, of aTc for the aTc switch, and 0.1 to 100 µM of cumate for the cumate switch (indicated in each experiment). The induction time is indicated in each experiment. The pSIP system induction was done using the IP673 peptide (GeneCust, Boyes, France), to a final concentration of 100 ng/mL.

### Flow cytometry analysis

Engineered *L. plantarum* WCSF1 was grown in 4 mL of MRS medium supplemented with 10 µg/mL erythromycin at 37 °C and 250 rpm. The next day, bacteria were sub-cultured to an OD_600nm_ of 0.01 in 2 mL of MRS medium supplemented with 10 µg/mL erythromycin and grown at 37 °C with shaking (250 rpm) for a period of 8 hours. For the induced conditions, inducers were added along with the erythromycin. After that, 1 mL of the bacterial cultures were pelleted down by centrifugation at 6000 rpm for 2 minutes. The supernatant was discarded carefully, and the pellets were resuspended in 1 mL of sterile Dulbecco’s 1X PBS (DPBS). Finally, the mixtures were serially diluted by a 10^4^-dilution factor, and 5,000 bacterial events were recorded for analysis using the Guava easyCyte BG flow-cytometer (Luminex, USA). The fluorescence intensity of mCherry was measured using excitation by a green laser at 532 nm (100 mW) and the Orange-G detection channel 620/52 nm filter was used for signal analysis. The Luminex GuavaSoft 4.0 software for EasyCyte was used for the analysis and representation of data.

### Microplate reader setup for induction levels assessment

Engineered *L. plantarum* WCSF1 was grown in 4 mL of MRS medium supplemented with 10 µg/mL erythromycin at 37 °C and 250 rpm. After overnight growth, bacteria were sub-cultured to low OD_600nm_ (indicated in the results section) in MRS medium supplemented with 10 µg/mL erythromycin. For the induced conditions, the inducers were added along with the erythromycin, and bacteria were incubated for a period of 5 to 8 hours (the exact time and cumate concentration are indicated in the results section). After that, bacterial pellets were resuspended in 1 mL of DPBS, and 200 µL of the mixture was transferred to a UV STAR Flat Bottom 96 well microtiter plate (Greiner BioOne GmbH, Germany). The microplate reader Infinite 200 Pro (Tecan Deutschland GmbH, Germany) was used to quantify the levels of fluorescence. The mCherry fluorescence intensity was measured using the excitation/emission values: Exλ / Emλ = 587 nm/625 nm. The fluorescence values were normalized with the OD_600nm_ of the mixtures (quantified by measuring the absorbance at 600 nm) to calculate the Relative Fluorescence Units (RFU) (Fluorescence/OD_600nm_). In all cases, the RFU values for *L. plantarum* WCFS1 carrying an empty vector (a plasmid based on the origin and the antibiotic resistance) were used as blank. The settings for the microplate reader were as follows: the Z-position was set at 19000 µm, the gain settings at 80-100 (indicated in the results section), and the readings were taken from the top.

### Microplate reader kinetics setup for induction assessment

Engineered *L. plantarum* WCSF1 was grown in 4 mL of MRS medium supplemented with 10 µg/mL erythromycin at 37 °C and 250 rpm. After overnight growth, bacteria were sub-cultured to low OD_600nm_ and incubated at 37 °C and 250 rpm.

- <u>In MRS medium</u>: after 8 hours of growth, bacteria were again diluted to an OD_600nm_ of 0.01 with MRS supplemented with 10 µg/mL erythromycin. For the induced conditions, 5 µM of cumate, 50 µM of vanillate, or 100 ng/mL of IP673 peptide were added for induction. Next, bacteria were transferred to a 96-well plate, and a kinetic cycle was set at 37°C or 30°C for up to 16 hours (indicated in the results section). Fluorescence intensity and OD_600nm_ values were taken every 30 minutes. RFU values were calculated as described in the previous section.
- <u>In LB or LBG media</u>: when bacteria reached an OD_600nm_ of 0.1 or 0.3, bacteria were transferred into either LB medium or LBG medium (LB supplemented with 0.1M of glucose) supplemented with 10 µg/mL erythromycin. For the induced conditions, 5 µM of cumate was added. Next, bacteria were transferred to a 96-well plate, and a kinetic cycle was set at 37°C for up to 21 hours. Fluorescence intensity and OD_600nm_ values were taken every 30 minutes. RFU values were calculated as described in the previous section.

### pH measurements

For non-encapsulated bacteria experiments, the pH of the media was measured after 0, 10, 15, and 20 hours (indicated in each experiment). For encapsulated bacteria experiments, PEARLs were kept for 1 to 3 days in the incubator. The pH of the LB medium was measured after every 24 hours. In all cases, the supernatants were collected and centrifuged for 3 minutes at 10000 rpm. The new supernatants were transferred into new tubes, and the pH was measured using a calibrated Elite pH Spear Pocket Tester (Thermo Scientific, Germany).

### NucA Assay (in DNase medium with Methyl green)

#### Qualitative Assay

Engineered *L. plantarum* WCSF1 was grown overnight in MRS medium supplemented with 10 µg/mL erythromycin at 37°C, 250 rpm. The next day, the OD_600nm_ was measured, and bacteria were subcultured in 4 mL of MRS medium supplemented with 10 µg/mL erythromycin at an OD_600nm_ of 0.05 and incubated for approximately 5 hours at 30 or 37°C until OD_600nm_ was close to 1. For the included conditions, 5 µM of cumate was added during the subculturing. 5 hours later, 10 µL of the cultures were spotted on DNase agar with Methyl green (per liter: 20 g of tryptose, 5 g of sodium chloride, 2 g of deoxyribonucleic acid, 0.05 g of methyl green, and 13 g of agar; Altmann Analytik GmbH, Germany) supplemented with 10 µg/mL of erythromycin and incubated at 37°C for 40 hours. After this incubation, the BioRad Gel Documentation System was used to capture images of the DNase agar plates, and the fluorescence of the halo generated around the spotted area was measured using ImageJ (version 1.53k).

#### Quantitative Assay

Engineered *L. plantarum* WCSF1 was grown as described in the qualitative assay section. When bacteria reached an OD_600nm_ of approximately 1, bacteria equivalent to OD_600nm_ of 0.1 were transferred to 1 mL of DNase medium supplemented with 10 µg/mL erythromycin and incubated at 30 or 37°C and 250 rpm shaking conditions for 24 hours. For the induced conditions, 5 µM of cumate was added during the subculturing. The DNase medium was obtained as described in [12]. After 24 hours, the cultures were centrifuged at 7000 rpm and 200 µL of the supernatants were added to the clear bottom 96-well microtiter plate. The samples were analyzed in the microplate reader Infinite 200 Pro was used to measure the fluorescence intensity of the supernatants (Exλ / Emλ = 633/668 nm). The Z-position was set at 19000 µm and the gain settings at 100. The NucA concentration was correlated and calculated using the standard curve at 24 hours described in [12].

#### Kinetics in the microplate reader

Engineered *L. plantarum* WCSF1 was grown in 4 mL of MRS medium supplemented with 10 µg/mL erythromycin at 37 °C and 250 rpm. After overnight growth, bacteria were sub-cultured to an OD_600nm_ of 0.01 in MRS medium supplemented with 10 µg/mL erythromycin. When bacteria reached an OD_600nm_ of approximately 0.1 and 0.3, bacteria were transferred to DNase medium with and without cumate and supplemented with 10 µg/mL erythromycin. 200 µL of the mixture was transferred to a 96-well plate and a kinetic study was set for 20 hours. NucA secretion was measured by taking measurements of the DNase medium at Ex/Em = 633/668nm and absorbance at 600 nm. Measurements were taken every 30 minutes. The manual gain was set at 100, Z-position at 19000 µm, the temperature was 37 °C, and the integration time was 20 µs.

### PEARLs fabrication and cumate-responsiveness

#### Fabrication

PEARLs were fabricated as described in [12]. In brief, 3 wt% alginate solution was prepared by dissolving alginate in sterile Milli-Q water, followed by autoclaving the solution for 15 minutes at 121°C. Equal amounts of 3 wt% alginate and *L. plantarum* WCSF1 were resuspended in LB medium supplemented with 10 µg/mL erythromycin (approximately 10^8^ cells per mL) were mixed and vortexed to obtain the core solution (1.5 wt%). Equal amounts of 3 wt% alginate and sterile Milli-Q water were mixed to obtain the shell solution (1.5 wt%). 10 µL droplets of core solution were pipetted and dropped into a 5 wt% CaCl2 solution (ionic crosslinker). The droplets were left to polymerize in the ionic crosslinker for 15 minutes. After that, the core beads were collected and washed thrice with sterile Milli-Q water. Alginate core beads were dipped into a 2-mL eppendorf with 1.5 wt% shell alginate solution and the core bead with 25 µL of shell solution was pipetted and dropped into a 5 wt% cross-linker solution. Core-shell PEARLs were left to polymerize in the cross-linker solution for 15 minutes before the induction experiments.

#### Cumate induction and quantification

PEARLs were transferred into a 96-well plate, and 200 µL of LB medium or Dulbecco’s Modified Eagle Medium (DMEM) (Sigma-Aldrich, Germany) supplemented with 10 µg/mL erythromycin were added. For the induced conditions, 100 µM of cumate or 50 µM of vanillate were added. Next, the microplate reader was used to track bacterial leakiness (bacterial growth as an increase in absorbance at 600 nm), and mCherry production (Ex/Em = 587/625 nm) for 24 to 70 hours depending on the experiments. Measurements were taken every 30 minutes. The manual gain was set at 100, the Z-position at 19000 µm, the temperature was 37 °C, and the integration time was 20 µs. The BioRad Gel Documentation System was also used to capture macroscopic images of the PEARLs after induction. Images were processed with ImageJ.

For the NucA experiments, 200 µL of DNase (supplemented with 10 µg/mL erythromycin) medium without or with cumate (100 µM) was added. NucA secretion was measured by taking measurements of the DNase medium surrounding the PEARLs at Ex/Em = 633/668nm and absorbance at 600 nm after 24 hours at 37°C. The manual gain was set at 100, the Z-position at 19000 µm, and the integration time was 20 µs. After that, the wells were washed thrice with PBS. Next, 200 µL of fresh DNase medium (supplemented with 10 µg/mL erythromycin) medium without or with cumate (100 µM) was added, and measurements were taken 24 hours after (day 2). The same procedure was repeated until day 7 or day 9 depending on the experiment.

#### Live/Dead staining and imaging of PEARLs

A sterile scalpel was used to carefully detach the shells from the cores of the PEARLS and to split the cores in half. Next, the half cores were stained with a mixture of SYTO9 and propidium iodide (PI) (Thermo Fisher Scientific) at final concentrations of 5 µM and 30 µM in PBS, respectively, for 30 minutes at room temperature in static. After that, the stained cores were washed with PBS before imaging.

Imaging was performed using a Zeiss Cell Discoverer 7 microscope equipped with Zen 3.11 software (Zeiss, Oberkochen, Germany). A Zeiss Plan-Apochromat 20×/0.95 objective with an optovar 0.5× Tubelens was used for image acquisition. Excitation/emission wavelengths were set to 488 nm/410–546 nm and 561 nm/605– 700 nm for detecting live and dead bacterial populations, respectively. A tile scan of the entire PEARL sample was acquired at a Z-stack interval of 5,52 µm, taken 50 µm above the plate surface. The 3D volumes of live and dead bacterial colonies were quantified using the Blob Finder tool (ZEISS Arivis Pro, Zeiss, Oberkochen, Germany). In addition to colonies stained exclusively with SYTO9 (live) or PI (dead), some colonies showed partial staining with both markers. For these SYTO9/PI-positive colonies, the volume was evenly divided between live and dead categories. The percentage of live cells (Live %) was calculated as the volume of live cells divided by the total volume of live and dead cells. Representative images were generated using orthographic maximum intensity projections (MIP) of the z-stacks (Zen 3.11, Zeiss, Oberkochen, Germany).

### Micro-PEARLs fabrication, secretion assessment and cumate responsiveness

#### Fabrication

The core and shell solutions were prepared in the same compositions as prepared for PEARLs. 5 mL of core solution was filled in a 5 mL syringe and 20 mL of shell solution was filled in a 20 mL syringe. Micro-PEARLs were fabricated using the microencapsulator B-390 (Buchi, Switzerland). The autoclaved reaction vessel was arranged with a 150 µm nozzle (for the core) and a 300 µm nozzle (for the shell). Syringes with core and shell alginate solutions were secured to the respective entry points of the aseptic reaction vessel. 200 mL of crosslinker solution (5 wt% CaCl2 solution) was introduced into the reaction vessel through the hose with a filter (Sartorius, Germany). The microencapsulator device was switched on and voltage (1600 V), frequency (910 Hz), syringe pump for core (20 mL/second), syringe pump for shell (7 mL/second) and a stirrer (40 %) were enabled. The micro-PEARLs with core and shell were dropped into the reaction vessel until the core solution ran out. The micro-PEARLs were left in the reaction vessel for up to 15 minutes for polymerization. The micro-PEARLs were collected in a sterile schott flask connected to the device through an efflux port at the bottom of the reaction vessel. The micro-PEARLs were finally washed thrice with sterile Milli-Q water. The reaction vessel and the nozzles were cleaned by pumping sterile water and the device was autoclaved at 121 °C.

#### NucA secretion assessment

After the washes, the micro-PEARLs were collected with a sterile spatula and placed in the center of DNase agar (Altmann Analytik GmbH, Germany) plates supplemented with 10 µg/mL of erythromycin and incubated at 37°C for 5 days. The next day, the discoloration zone was captured using the BioRad Gel Documentation System. The images were processed using ImageJ.

#### Cumate induction

After three washes with sterile Milli-Q water, the micro-PEARLs were collected with a sterile spatula and placed in either 2-mL tubes or a 12-well plate. For the induction in the tubes, 1 mL of LB medium supplemented with 10 µg/mL of erythromycin and 100 µM of cumate was added. Induction was performed for approximately 24 hours. After that, induction was assessed with the BioRad Gel Documentation System or the Keyence microscope (BZ-X800E). For induction in the plates, 500 µL of LB supplemented with 10 µg/mL of erythromycin and 100 µM of cumate was added per well. The plate was placed in the microplate reader and an increase in mCherry production (Ex/Em = 587/625 nm) was tracked for 19 hours approximately. The absorbance at 600 nm was also measured for normalization of the values. Measurements were taken every 30 minutes. The manual gain was set at 100, Z-position at 23000 µm, the temperature was 37 °C, and the integration time was 20 µs.

#### Fluorescence Microscopy

The Keyence microscope (BZ-X800E) was used to perform microscopy of micro-PEARLs. After the induction, micro-PEARLs were transferred to a 12-well plate with a glass bottom for better imaging. Multichannel images of micro-PEARLs were captured in 4X and 10X (S plan fluor objective) magnification using the CH3 (red fluorescence) and CH4 (brightfield) channels. The images were processed and analyzed using ImageJ.

### Statistical analysis

Statistical analysis was done using GraphPad Prism 7.0 software. Student’s t-tests were used to determine statistically significant differences among the means of the groups. The differences among groups are indicated as: no statistically significant difference (ns), *p*-values <0.05 (*), *p*-values <0.01 (**), *p*-values <0.005 (***), *p*-values <0.001 (****).

## SUPPORTING INFORMATION

Supporting information is available from the Wiley Online Library.

## FUNDIN1G

This work was supported by a Deutsche Forschungsgemeinschaft (DFG) Research grant (Project # 455063657), a DFG Collective Research Center (SFB1027) subproject grant (Project #200049484), and the Leibniz Association through the Leibniz Science Campus on Living Therapeutic Materials (LifeMat).

## DATA AVAILABILITY

The raw and processed datasets along with relevant metadata are available from the corresponding author upon reasonable request.

## Supporting information

Supporting Information_revised

## ACKNOWLEDGEMENTS

We thank Prof. Geir Mathiesen (Norwegian University of Life Sciences) for sharing the pLp_3050sNuc plasmid (Addgene plasmid # 122030) with us. The *L. plantarum* WCFS1 strain was a kind gift from Prof. Gregor Fuhrmann (Friedrich-Alexander-Universität, Erlangen-Nürnberg). The authors used Biorender for the illustrations of this study. We thank the group of Prof. Wilfried Weber for granting access to their Buchi B-390 microencapsulator device for the fabircation of the micro-PEARLs and providing initial training for the device.

## REFERENCES

1. Giraffa, G.; Chanishvili, N.; Widyastuti, Y., Research in Microbiology 2010, 161 (6), 480–487. DOI 10.1016/j.resmic.2010.03.001.

2. Zhang, Z.; Lv, J.; Pan, L.; Zhang, Y., Applied Microbiology and Biotechnology 2018, 102 (19), 8135–8143. DOI 10.1007/s00253-018-9217-9.

3. Monachese, M.; Burton Jeremy, P.; Reid, G., Applied and Environmental Microbiology 2012, 78 (18), 6397–6404. DOI 10.1128/AEM.01665-12.

4. O’Callaghan, J.; O’Toole, P. W., Lactobacillus: Host–Microbe Relationships. In Between Pathogenicity and Commensalism, Dobrindt, U.; Hacker, J. H.; Svanborg, C., Eds. Springer Berlin Heidelberg: Berlin, Heidelberg, 2013; pp 119–154.

5. Chee, W. J. Y.; Chew, S. Y.; Than, L. T. L., Microbial Cell Factories 2020, 19 (1), 203. DOI 10.1186/s12934-020-01464-4.

6. Wang, J.; Zhang, H.; Chen, X.; Chen, Y.; Menghebilige; Bao, Q., Journal of Dairy Science 2012, 95 (4), 1645–1654. DOI 10.3168/jds.2011-4768.

7. Santos Rocha, C.; Lakhdari, O.; Blottière, H. M.; Blugeon, S.; Sokol, H.; Bermu’dez-Humara’n, L. G.; Azevedo, V.; Miyoshi, A.; Doré, J.; Langella, P.; Maguin, E.; van de Guchte, M., Inflammatory Bowel Diseases 2012, 18 (4), 657–666. DOI 10.1002/ibd.21834.

8. Zhang, Z.; Niu, H.; Qu, Q.; Guo, D.; Wan, X.; Yang, Q.; Mo, Z.; Tan, S.; Xiang, Q.; Tian, X.; Yang, H.; Liu, Z., Critical Reviews in Food Science and Nutrition, 1–22. DOI 10.1080/10408398.2024.2448562.

9. Blanch-Asensio, M.; Dey, S.; Tadimarri, V. S.; Sankaran, S., Microbial Biotechnology 2024, 17 (1), e14335. DOI 10.1111/1751-7915.14335.

10. Scheppler, L.; Vogel, M.; Zuercher, A. W.; Zuercher, M.; Germond, J.-E.; Miescher, S. M.; Stadler, B. M., Vaccine 2002, 20 (23), 2913–2920. DOI 10.1016/S0264-410X(02)00229-3.

11. Pan, N.; Liu, B.; Bao, X.; Zhang, H.; Sheng, S.; Liang, Y.; Pan, H.; Wang, X., Vaccines 2021, 9 (9), 984.

12. Tadimarri, V. S.; Blanch-Asensio, M.; Deshpande, K.; Baumann, J.; Baumann, C.; Müller, R.; Trujillo, S.; Sankaran, S., Small 2025, n/a (n/a), 2408316. DOI 10.1002/smll.202408316.

13. Meng, Q.; Yuan, Y.; Li, Y.; Wu, S.; Shi, K.; Liu, S., ACS Synthetic Biology 2021, 10 (7), 1728–1738. DOI 10.1021/acssynbio.1c00123.

14. Peirotén, Á.; Landete, J. M., Applied Microbiology and Biotechnology 2020, 104 (9), 3797–3805. DOI 10.1007/s00253-020-10426-0.

15. Rud, I.; Jensen, P. R.; Naterstad, K.; Axelsson, L., Microbiology 2006, 152 (4), 1011–1019. DOI 10.1099/mic.0.28599-0.

16. Heiss, S.; Hörmann, A.; Tauer, C.; Sonnleitner, M.; Egger, E.; Grabherr, R.; Heinl, S., Microbial Cell Factories 2016, 15 (1), 50. DOI 10.1186/s12934-016-0448-0.

17. Zhang, S.; Xu, Z.; Qin, L.; Kong, J., Biochemical Engineering Journal 2019, 151, 107316. DOI 10.1016/j.bej.2019.107316.

18. Martínez-Fernández, J. A.; Bravo, D.; Peirotén, Á.; Arqués, J. L.; Landete, J. M., Applied Microbiology and Biotechnology 2019, 103 (9), 3819–3827. DOI 10.1007/s00253-019-09743-w.

19. Karlskås, I. L.; Maudal, K.; Axelsson, L.; Rud, I.; Eijsink, V. G. H.; Mathiesen, G., PLOS ONE 2014, 9 (3), e91125. DOI 10.1371/journal.pone.0091125.

20. Sørvig, E.; Mathiesen, G.; Naterstad, K.; Eijsink, V. G. H.; Axelsson, L., Microbiology 2005, 151 (7), 2439–2449. DOI 10.1099/mic.0.28084-0.

21. Nguyen, T.-T.; Nguyen, T.-H.; Maischberger, T.; Schmelzer, P.; Mathiesen, G.; Eijsink, V. G. H.; Haltrich, D.; Peterbauer, C. K., Microbial Cell Factories 2011, 10 (1), 46. DOI 10.1186/1475-2859-10-46.

22. Yang, W.-T.; Li, Q.-Y.; Ata, E. B.; Jiang, Y.-L.; Huang, H.-B.; Shi, C.-W.; Wang, J.-Z.; Wang, G.; Kang, Y.-H.; Liu, J.; Yang, G.-L.; Wang, C.-F., Applied Microbiology and Biotechnology 2018, 102 (19), 8307–8318. DOI 10.1007/s00253-018-9238-4.

23. Halbmayr, E.; Mathiesen, G.; Nguyen, T.-H.; Maischberger, T.; Peterbauer, C. K.; Eijsink, V. G. H.; Haltrich, D., Journal of Agricultural and Food Chemistry 2008, 56 (12), 4710–4719. DOI 10.1021/jf073260+.

24. Meyer, A. J.; Segall-Shapiro, T. H.; Glassey, E.; Zhang, J.; Voigt, C. A., Nature Chemical Biology 2019, 15 (2), 196–204. DOI 10.1038/s41589-018-0168-3.

25. Brautaset, T.; Lale, R.; Valla, S., Microbial Biotechnology 2009, 2 (1), 15–30. DOI 10.1111/j.1751-7915.2008.00048.x.

26. Mao, J.; Zhang, H.; Chen, Y.; Wei, L.; Liu, J.; Nielsen, J.; Chen, Y.; Xu, N., Biotechnology Advances 2024, 74, 108401. DOI 10.1016/j.biotechadv.2024.108401.

27. Yoon, S.; Seo, K. S.; Park, N.; Kim, C.; Dey, P.; Thornton, J. A.; Park, J. Y., Scientific Reports 2024, 14 (1), 30780. DOI 10.1038/s41598-024-81001-0.

28. Wang, Y.; Liu, Y.; Li, J.; Chen, Y.; Liu, S.; Zhong, C., Current Opinion in Chemical Biology 2022, 70, 102188. DOI 10.1016/j.cbpa.2022.102188.

29. Blanch-Asensio, M.; Tadimarri, V. S.; Wilk, A.; Sankaran, S., Microbial Cell Factories 2024, 23 (1), 42. DOI 10.1186/s12934-024-02302-7.

30. Bina, X. R.; Wong, E. A.; Bina, T. F.; Bina, J. E., Plasmid 2014, 76, 87–94. DOI 10.1016/j.plasmid.2014.10.004.

31. Zhang, L.; Zou, W.; Ni, M.; Hu, Q.; Zhao, L.; Liao, X.; Huang, Q.; Zhou, R., Microbiology Spectrum 2022, 10 (4), e00363–22. DOI 10.1128/spectrum.00363-22.

32. Lutz, R.; Bujard, H., Nucleic Acids Research 1997, 25 (6), 1203–1210. DOI 10.1093/nar/25.6.1203.

33. Klotz, A.; Kaczmarczyk, A.; Jenal, U., Applied and Environmental Microbiology 2023, 89 (6), e00211–23. DOI 10.1128/aem.00211-23.

34. Seo, S.-O.; Schmidt-Dannert, C., Applied Microbiology and Biotechnology 2019, 103 (1), 303–313. DOI 10.1007/s00253-018-9485-4.

35. Kunjapur, A. M.; Prather, K. L. J., ACS Synthetic Biology 2019, 8 (9), 1958–1967. DOI 10.1021/acssynbio.9b00071.

36. Tan, C.; Marguet, P.; You, L., Nature Chemical Biology 2009, 5 (11), 842–848. DOI 10.1038/nchembio.218.

37. Daniel, R.; Rubens, J. R.; Sarpeshkar, R.; Lu, T. K., Nature 2013, 497 (7451), 619–623. DOI 10.1038/nature12148.

38. Dey, S.; Blanch-Asensio, M.; Balaji Kuttae, S.; Sankaran, S., Microbial Biotechnology 2023, 16 (6), 1264–1276. DOI 10.1111/1751-7915.14228.

39. Dri, A.-M.; Moreau, P. L., Molecular Microbiology 1994, 12 (4), 621–629. DOI 10.1111/j.1365-2958.1994.tb01049.x.

40. Krukenberg, K. A.; Southworth, D. R.; Street, T. O.; Agard, D. A., Journal of Molecular Biology 2009, 390 (2), 278–291. DOI 10.1016/j.jmb.2009.04.080.

41. Prost, L. R.; Daley, M. E.; Le Sage, V.; Bader, M. W.; Le Moual, H.; Klevit, R. E.; Miller, S. I., Molecular Cell 2007, 26 (2), 165–174. DOI 10.1016/j.molcel.2007.03.008.

42. Ramos, C. L.; Thorsen, L.; Ryssel, M.; Nielsen, D. S.; Siegumfeldt, H.; Schwan, R. F.; Jespersen, L., Research in Microbiology 2014, 165 (3), 215–225. DOI 10.1016/j.resmic.2014.02.005.

43. Bai, F.; Li, Z.; Umezawa, A.; Terada, N.; Jin, S., Biotechnology Advances 2018, 36 (2), 482–493. DOI 10.1016/j.biotechadv.2018.01.016.

44. Neef, J.; van Dijl, J. M.; Buist, G., Essays in Biochemistry 2021, 65 (2), 187–195. DOI 10.1042/EBC20200171.

45. Reed, B.; Chen, R., Research in Microbiology 2013, 164 (6), 675–682. DOI 10.1016/j.resmic.2013.03.006.

46. Mathiesen, G.; Sveen, A.; Brurberg, M. B.; Fredriksen, L.; Axelsson, L.; Eijsink, V. G. H., BMC Genomics 2009, 10 (1), 425. DOI 10.1186/1471-2164-10-425.

47. Le Loir, Y.; Nouaille, S.; Commissaire, J.; Brétigny, L.; Gruss, A.; Langella, P., Applied and Environmental Microbiology 2001, 67 (9), 4119–4127. DOI 10.1128/AEM.67.9.4119-4127.2001.

48. Nouaille, S.; Morello, E.; Cortez-Peres, N.; Le Loir, Y.; Commissaire, J.; Gratadoux, J. J.; Poumerol, E.; Gruss, A.; Langella, P., Applied and Environmental Microbiology 2006, 72 (3), 2272–2279. DOI 10.1128/AEM.72.3.2272-2279.2006.

49. Birnbaum, D. P.; Manjula-Basavanna, A.; Kan, A.; Tardy, B. L.; Joshi, N. S., Advanced Science 2021, 8 (11), 2004699. DOI 10.1002/advs.202004699.

50. Gilbert, C.; Tang, T.-C.; Ott, W.; Dorr, B. A.; Shaw, W. M.; Sun, G. L.; Lu, T. K.; Ellis, T., Nature Materials 2021, 20 (5), 691–700. DOI 10.1038/s41563-020-00857-5.

51. Johnston, T. G.; Yuan, S.-F.; Wagner, J. M.; Yi, X.; Saha, A.; Smith, P.; Nelson, A.; Alper, H. S., Nature Communications 2020, 11 (1), 563. DOI 10.1038/s41467-020-14371-4.

52. Rojas-Muñoz, Y. V.; de Jesús Perea-Flores, M.; Quintanilla-Carvajal, M. X. Probiotic Encapsulation: Bead Design Improves Bacterial Performance during In Vitro Digestion (Part 2: Operational Conditions of Vibrational Technology) Polymers [Online], 2024.

53. Berninger, T.; Mitter, B.; Preininger, C., Journal of Microencapsulation 2016, 33 (2), 127–136. DOI 10.3109/02652048.2015.1134690.

54. Lyu, Y.; Huang, H.; Su, Y.; Ying, B.; Liu, W.-C.; Dong, K.; Du, N.; Langer, R. S.; Gu, Z.; Nan, K., Matter 2024, 7 (4), 1440–1465. DOI 10.1016/j.matt.2024.01.031.

55. Harimoto, T.; Hahn, J.; Chen, Y.-Y.; Im, J.; Zhang, J.; Hou, N.; Li, F.; Coker, C.; Gray, K.; Harr, N.; Chowdhury, S.; Pu, K.; Nimura, C.; Arpaia, N.; Leong, K. W.; Danino, T., Nature Biotechnology 2022, 40 (8), 1259–1269. DOI 10.1038/s41587-022-01244-y.

56. Altin-Yavuzarslan, G.; Drake, K.; Yuan, S.-F.; Brooks, S. M.; Kwa, E.; Alper, H. S.; Nelson, A., Matter 2025, 8 (1). DOI 10.1016/j.matt.2024.10.008.

57. Blanch-Asensio, M.; Dey, S.; Sankaran, S., PLOS ONE 2023, 18 (2), e0281625. DOI 10.1371/journal.pone.0281625.

