## Supporting Information_revised for "Encapsulation-enhanced genetic switches in lactobacilli"

| Primer name | Sequence |
|---|---|
| **Vector switch fw** | 5'- atggtttcaaagggtgaa -3' |
| **Vector switch rev** | 5'- tctatctatatctctagcgag -3' |
| **Repressor removal fw** | 5'- gatcgttcgtagagatcg -3' |
| **Repressor removal rev** | 5'- tctatctatatctctagcgag -3' |
| **Operon CymR vector fw** | 5'- aataactagcataacccctt -3' |
| **Operon CymR vector rev** | 5'- ttacttgtataattcatccatacc -3' |
| **Operon CymR insert fw** | 5'- tggatgaattatacaagtaatagaattaaattaaaaagggagg -3' |
| **Operon CymR insert rev** | 5'- aaggggttatgctagttattttagcgtttgaatttagc -3' |
| **Promoter swap vector fw** | 5'- ccaaatataatgtctccaaa -3' |
| **Promoter swap vector rev** | 5'- tctatctatatctctagcgag -3' |
| **Promoter swap $P_{tec}$ fw** | 5'- tcgctagagatatagatagag -3' |
| **Promoter swap $P_{tec}$ rev** | 5'- tttggagacattatatttggtctcctttcgttcttgat -3' |
| **Promoter swap $P_{tlpA}$ fw** | 5'- tcgctagagatatagatagaggcttcaatagagttcttag -3' |
| **Promoter swap $P_{tlpA}$ rev** | 5'- tttggagacattatatttggcctccttcttaaagttaaac -3' |
| **NucA CymR vector fw** | 5'- gtccgagactaattcatg -3' |
| **NucA CymR vector rev** | 5'- ttcgttttctcctttcgt -3' |

**Table S1**: List of primers used in this study.

- **aTc genetic switch eBlock**

tcgctagagatatagatagataagcgttcggtcagtaaataatagaaataaaaaatcagacctaagactgatgacaaaaag
agaaaattttgataaaatagtcttagaattaaattaaaaagggaggccaaatata**atgagtcgtcttgacaaaagtaaagtca
tcaattcagccttagaacttcttaacgaggtaggcattgagggtcttactactagaaaattagcacagaagttaggcgtagaa
caacctacattatattggcatgtaaagaacaaacgtgcattattggatgcccttgcaatcgagatgcttgatcgtcaccacac
tcatttttgccctttggaaggagagagttggcaagattttcttcgcaataacgctaaatcatttcgatgtgctcttctttctcaccg
tgatggagcaaaagttcatttgggaacccgaccaacagagaagcaatacgaaacattagaaaatcagcttgcatttttatgc
caacagggattctctcttgagaatgcactttacgctcttagtgcagtcggccatttcacattgggatgtgtattggaggaccag
gagcaccaggtcgctaaagaagaacgtgagacccctaccaccgactcaatgccaccacttcttcgccaagcaatcgagtt
attcgatcatcaaggtgccgagcctgcattccttttcggccttgaacttatcatttgcggccttgaaaaacaattaaaatgtga
gtctggcagt**taaaataaggctcaccttcgggtggcctttctgcgctgcattaagcttttaaccacaagttgagcaagagat
cgttcgtagagatcgattgtttattttgtgtcattttgtttgacatccctatcagtgatagagatttgtgttgactcccggtctagaa
cggggtattattaaatccctatcagtgatagagagcatcaagaacgaaaggagaaaacgaa**atggtttcaaagggtgaag
a**

- **Cumate genetic switch eBlock (mCherry)**

tcgctagagatatagatagataagcgttcggtcagtaaataatagaaataaaaaatcagacctaagactgatgacaaaaag
agaaaattttgataaaatagtcttagaattaaattaaaaagggaggccaaatata**atgtctccaaaacgccgcacccaggc**

**agaacgagctatggagacccagggcaaattgattgcagccgctttgggtgtcttacgagagaaggatacgcaggctttc**
**gaattgcagacgtaccaggcgccgccggagtttcacgaggtgcccagagtcaccattttccaacaaagcttgaacttttact**
**tgcaacattcgagtggttgtatgaacaaatcaccgaacgctcacgtgcccgccttgcaaaattaaaaccagaagatgatgt**
**cattcagcaaatgcttgatgacgctgctgagtttttttagatgacgatttctcaattggcttggatttaatcgtagcagccgacc**
**gtgatcctgcccttcgagaaggaatccaacgtactgtcgagcgcaaccgattcgtagttgaagatatgtggcttggagtatt**
**ggtaagtcgtggtttatcacgtgatgatgcagaagatattctttggttgatttttaatagtgtacgtggccttgttgtcagatcatt**
**gtggcaaaaagataaggaaagattcgagcgtgtccgcaactctactttagaaattgcaagagaacgctacgctaaattcaa**
**acgc**taaaataaggctcaccttcgggtgggcctttctgcgctgcattaagcttttaaccacaagttgagcaagagatcgttcg
tagagatcgaaaacaaacagacaatctggtctgtttgtagaatagttgctcggaatttgagcaaacaaacagacaatctggt
ctgtttgtatttgtgttgactcccggtctagaacggggtattattaaaaacaaacagacaatctggtctgtttgtagcatcaaga
acgaaaggagaaaacgaa**atggtttcaaagggtgaaga**

- **Vanillate genetic switch eBlock**

tcgctagagatatagatagataagcgttcggtcagtaaataatagaaataaaaaatcagacctaagactgatgacaaaaag
agaaaattttgataaaatagtcttagaattaaattaaaaagggaggccaaatata**atggacatgcctcgtattaaaccgggtc**
**agcgtgttatgatggcactgcgtaaaatgattgcaagcggtgaaatcaaaagtggtgaacgtattgcagaaattccgaccg**
**cagcagcactgggtgttagccgtatgccggttcgtatcgcactgcgttcactggaacaagaaggtctggttgttcgtctgggt**
**gcacgtggttatgcagcccgtggtgttagcagcgatcagattcgtgatgcaattgaagttcgtggtgttctggaaggttttgc**
**agcacgtcgtctggcagaacgtggtatgaccgcagaaacccatgcacgttttgttgtactgattgcagaaggtgaagcact**
**gtttgcagccggtcgcctgaatggtgaagatctggatcgttatgccgcatataatcaggcatttcatgataccctggttagcg**
**cagcaggtaatggtgcagttgaaagcgcactggcacgtaatggttttgaaccgtttgcagcagccggtgcactggccctgg**
**atctgatggacctgtctgccgaatatgaacatctgctggcagcacatcgtcagcatcaggcagttctggatgcagttagctgt**
**ggtgatgccgaaggtgcagaacgtattatgcgtgatcatgcactggcagcaattcgtaatgcaaaagtttttgaagcagcag**
**caagcgcaggcgcaccgctgggtgcagcatggtcaattcgtgcagattga**taaaataaggctcaccttcgggtgggccttt
ctgcgctgcattaagcttttaaccacaagttgagcaagagatcgttcgtagagatcgaagaatagttgctcggaatttgagca
attggatccaatttgtgttgactcccggtctagaacggggtattattaaagcatcattggatccaataagaacgaaaggaga
aaacgaa**atggtttcaaagggtgaaga**

- **NucA genetic switch eBlock**

Gaacgaaaggagaaaacgaaatgaaaaaatttaactttaaaaccatgttgctattagttttggctagttgtgtcttcggggtcg
tcgttaacgtgactactagtcttggaccacaaaccgcaatcaccgcccaggcctccaagaagctccatcatcaccatcatc
acccattg**tcaactaaaaaattacataaagaacctgcgactttaattaaagcgattgatggtgatacggttaaattaatgtaca**
**aaggtcaaccaatgacattcagactattattggttgatacacctgaaacaaagcatcctaaaaaaggtgtagagaaatatgg**
**tcctgaagcaagtgcatttacgaaaaaaatggtagaaaatgcaaagaaaattgaagtcgagtttgacaaaggtcaaagaa**
**ctgataaatatggacgtggcttagcgtatatttatgctgatggaaaaatggtaaacgaagctttagttcgtcaaggcttggcta**
**aagttgcttatgtttacaaacctaacaatacacatgaacaacatttaagaaaaagtgaagcacaagcgaaaaaagagaaat**
**taaatatttggagcgaagacaacgctgattcaggtcaa**taaaataactagcataaccccttggggcctctaaacgggtcttga
ggggtttttgctgaaaggaggaactatatccgaacgatcctctcagtgcgagtcgtccgagactaattcatgac

**Figure S0:** List of synthetic genes (eBlocks) used in this study. The start of the mCherry gene is indicated in bold and red. The $P_{23}$ and $P_{tec}$ promoters are shown underlined and in green and orange, respectively. The tetR, cymR and vanR genes are shown in bold and in yellow, blue and purple, respectively. The tetR, cymR and vanR operator sequences are underlined and in yellow, blue and purple, respectively. NucA coding sequence is shown in bold and green.

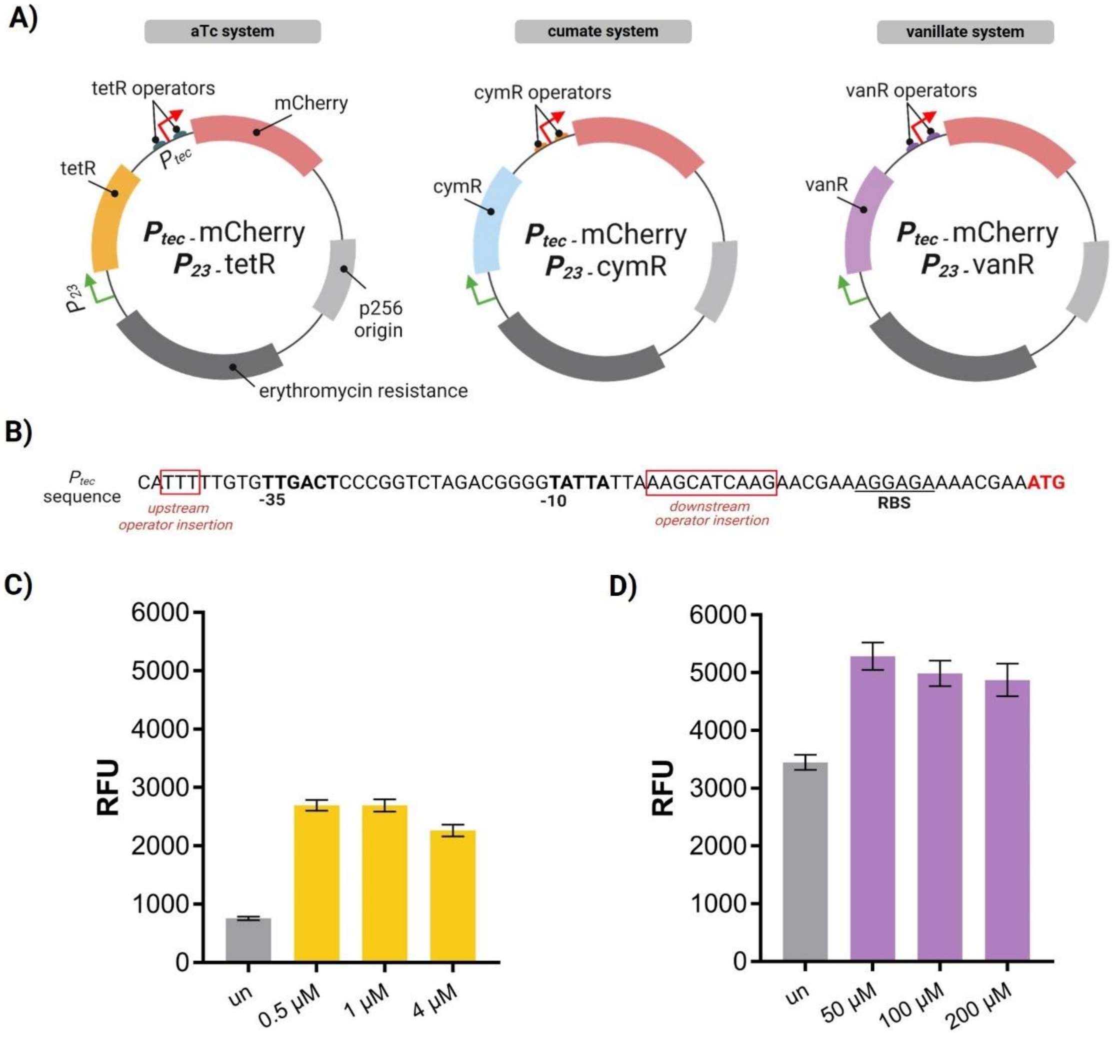


**Figure S1**: **A**) Plasmid maps for the aTc, cumate and vanillate switches. All the relevant genetic parts are depicted. **B**) Insertion of the operator sequences into $P_{tec}$ promoter. The upstream and downstream operators were inserted within the areas marked in red. **C**) RFU values after inducing with different aTc concentrations for 6 hours (starting $OD_{600nm}$ = 0.05, n = 1). **D**) RFU after inducing with different vanillate concentrations for 6 hours (starting $OD_{600nm}$ = 0.05, n = 1).

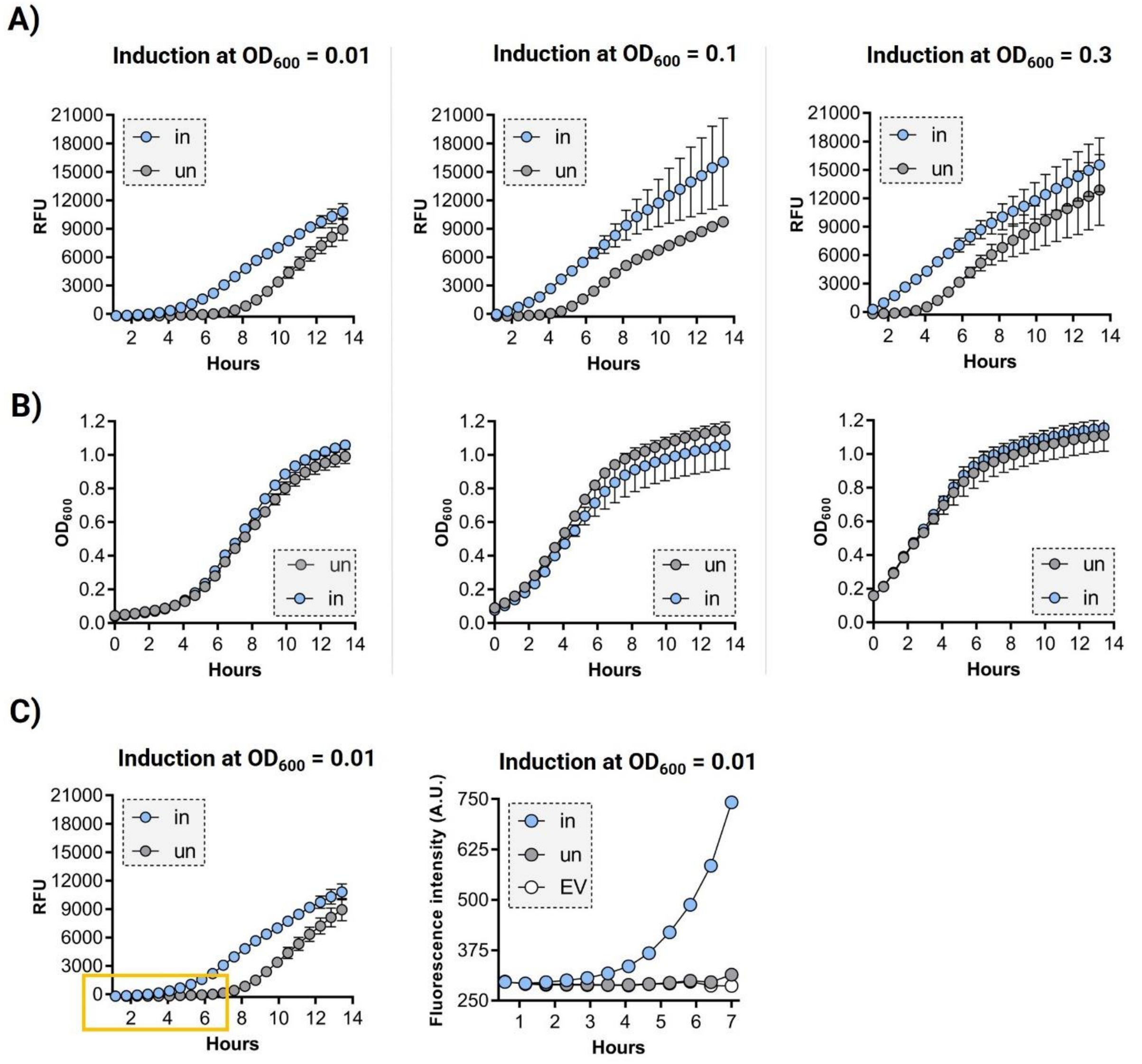


**Figure S2**: **A**) Kinetic plots showing the RFU values of the $P_{23}$_cymR variant when induced at different starting $OD_{600nm}$ (0.3, 0.1 and 0.01) for 14 hours at 37 °C (n = 3, mean ± SD). **B**) Plots showing the kinetic growth (as $OD_{600nm}$) of the $P_{23}$_cymR variant (uninduced and induced) at different starting $OD_{600nm}$ (0.3, 0.1 and 0.01) for 14 hours at 37 °C (n = 3, mean ± SD). "un" = uninduced condition, "in" = induced condition (5 µM of cumate). **C**) The left plot is the same as shown in panel A. The right plot is based on the fluorescence intensity values of the yellow square in the right plot. The EV values are not subtracted from the "un" (uninduced) and "in" (induced) conditions. Induction of the induced conditions started after 2 hours of growth, whereas the leakiness for the uninduced conditions started after 6.5 hours of growth (n = 3, mean ± SD).

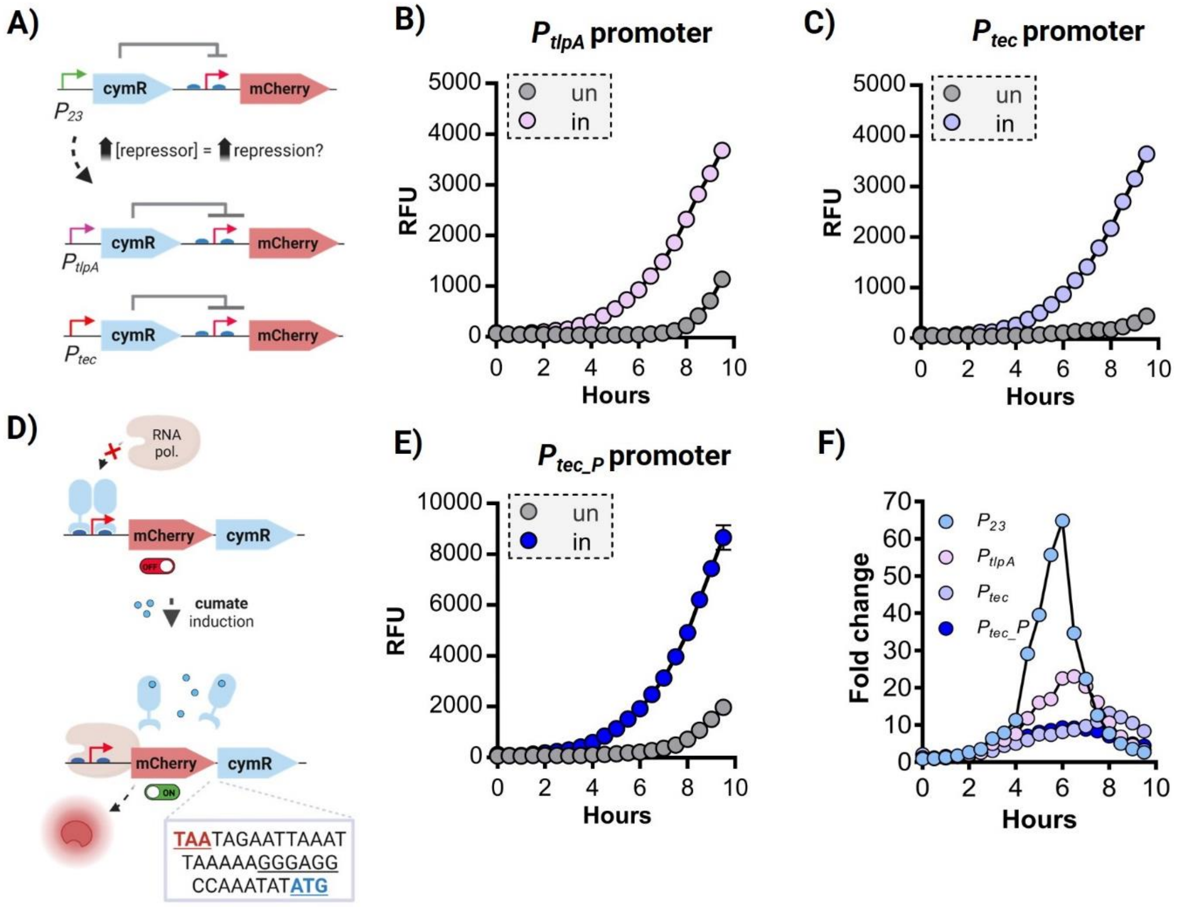


**Figure S3**: **A**) Scheme showing the $P_{23}$ promoter exchange for $P_{tlpA}$ and $P_{tec}$, thus generating the $P_{tlpA}$_cymR and the $P_{tec}$_cymR variants. **B**) RFU values for the $P_{tlpA}$_cymR variant after induction for ~10 hours (starting $OD_{600nm}$ = 0.01, n = 3, mean ± SD). "un" = uninduced condition, "in" = induced condition (5 µM of cumate). **C**) RFU values for the $P_{tec}$_cymR variant after induction for ~10 hours (starting $OD_{600nm}$ = 0.01, n = 3, mean ± SD). "un" = uninduced condition, "in" = induced condition (5 µM of cumate). **D**) Scheme showing the feedback circuit approach based on cloning the CymR repressor after the mCherry gene, both driven by the same (operated) promoter, $P_{tec}$ ($P_{tec}$_cymR_P variant). The exact DNA sequence between the mCherry STOP codon (highlighted in red) and the cymR ATG codon (highlighted in blue) is shown at the bottom. An additional RBS (the one used for the $P_{23}$ promoter) was included to ensure translation of the downstream gene (underlined). **E**) RFU values for the $P_{tec}$_cymR_P variant after induction for ~10 hours (starting $OD_{600nm}$ = 0.01, n = 3, mean ± SD). "un" = uninduced condition, "in" = induced condition (5 µM of cumate). **F**) Fold changes for all the cymR variants when grown at 37°C and induced and compared to 37 °C and uninduced.

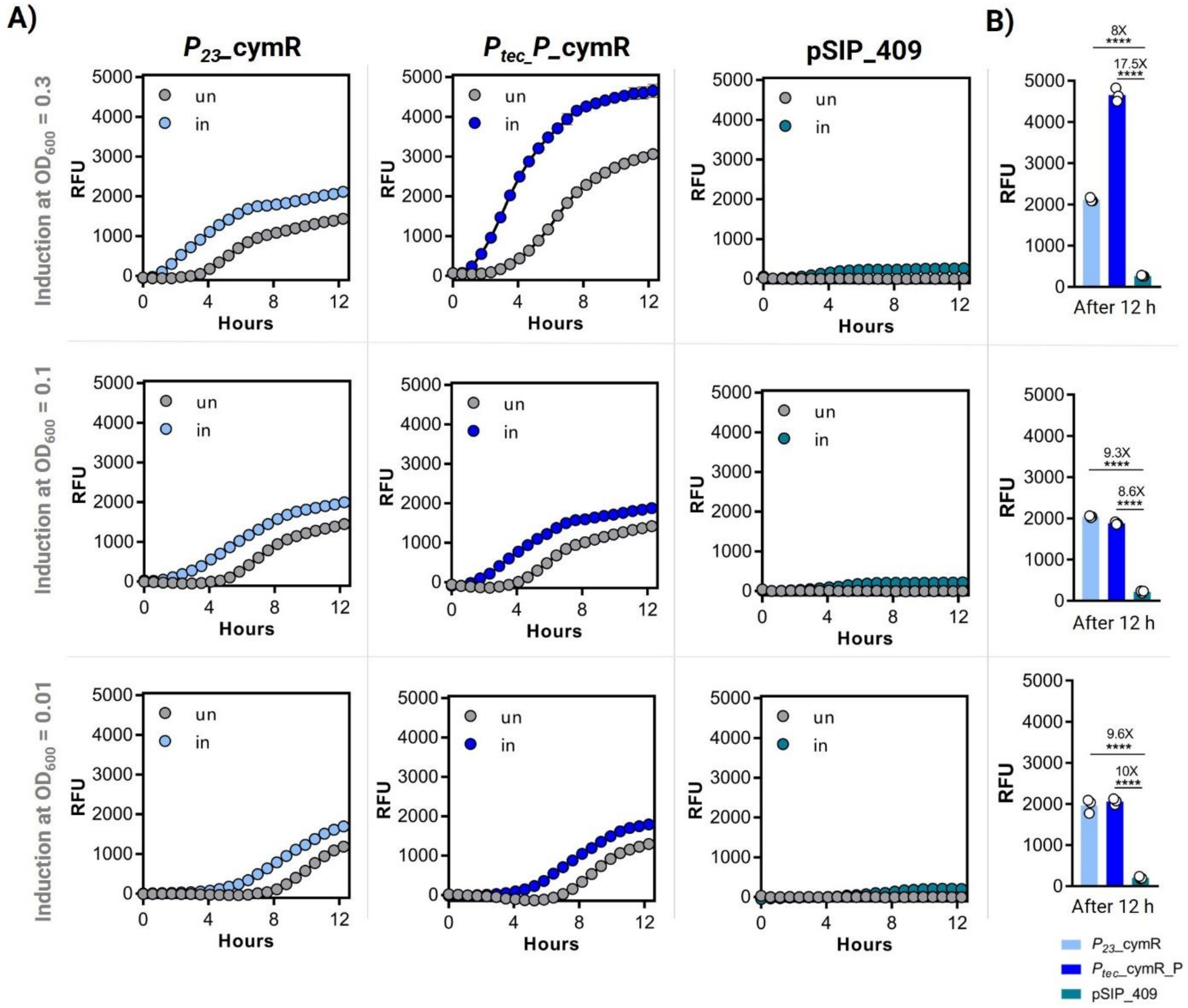


**Figure S4**: **A**) Comparison between the $P_{23}$_cymR, the $P_{tec}$_cymR_P, and the pSIP_409 variant. Bacteria were induced at three different $OD_{600nm}$ (0.01, 0.1 and 0.3) with 5 µM of cumate (for the cumate variants) and 100 ng/mL of IP673 (for the pSIP system) (n = 3, mean ± SD). "un" = uninduced condition, "in" = induced condition (5 µM of cumate). **B**) RFU values corresponding to the overall expression after 12 hours of induction for all three genetic switches (n = 3, mean ± SD). In these experiments, the microplate reader gain was set at 85 instead of 100 to avoid oversaturation of the detector by the mCherry expression of the $P_{tec}$_cymR_P variant.

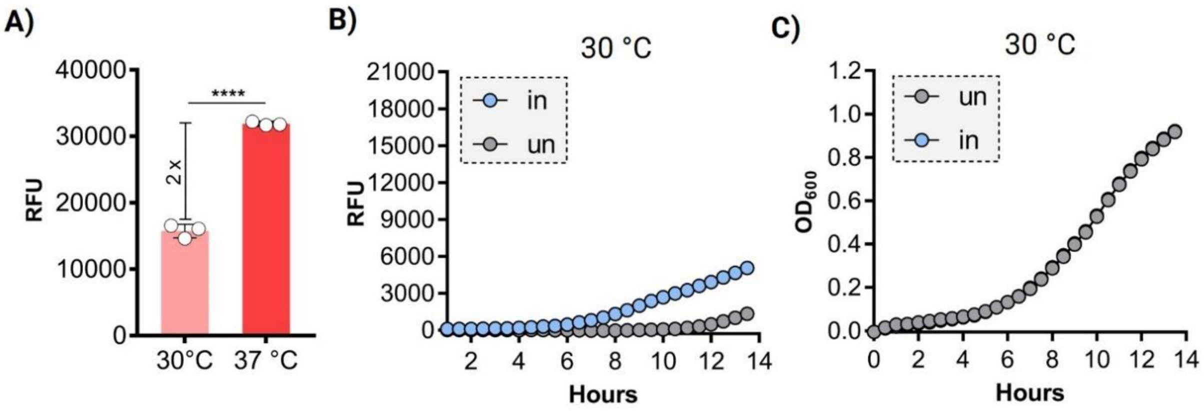


**Figure S5: A**) Thermo-responsiveness in terms of RFU for the $P_{tec}$ promoter when grown at 30 and 37°C for 16 hours (n = 3, mean ± SD). **B**) Kinetic plot showing the RFU values of the $P_{23}$_cymR variant (uninduced and induced) when grown at 30 °C (starting $OD_{600nm}$ = 0.01, n = 3, mean ± SD). **C**) Plot showing the kinetic growth (as $OD_{600nm}$) of the $P_{23}$_cymR variant (uninduced and induced) when grown at 30 °C (starting $OD_{600nm}$ = 0.01, n = 3, mean ± SD). “un” = uninduced condition, “in” = induced condition (5 μM of cumate).

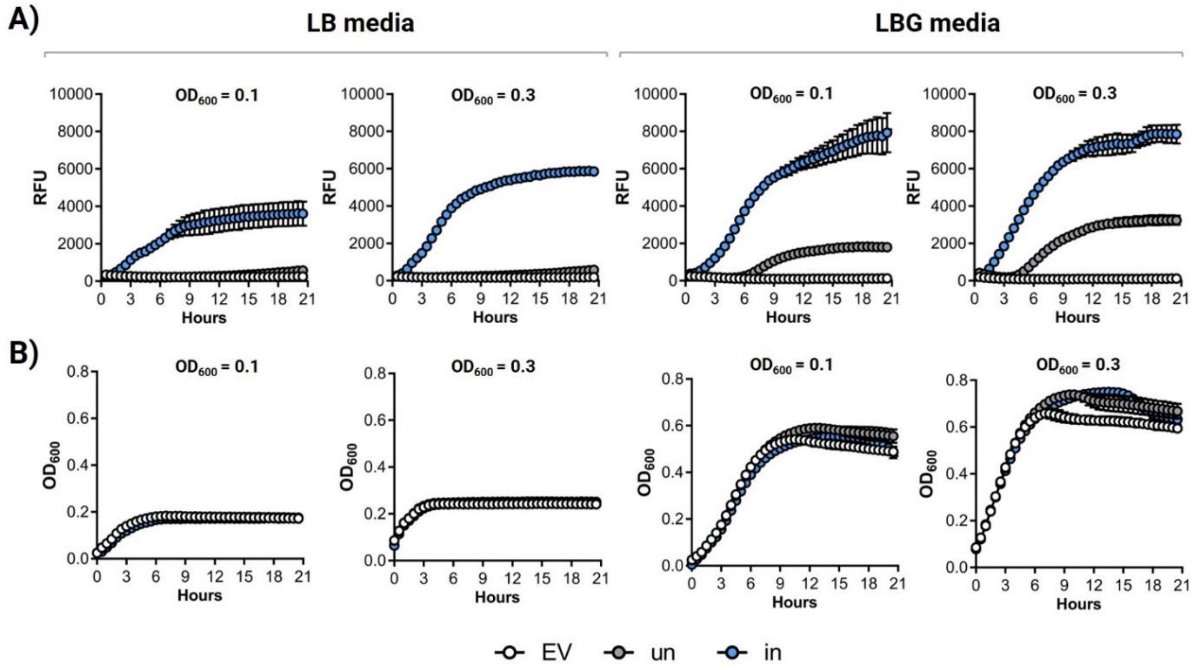


**C)**

| Media | pH (0h) | pH (10h) | pH (15h) | pH (20h) |
|---|---|---|---|---|
| LB 0.1 | 6.79 | 5.93 | 5.86 | 5.74 |
| LB 0.3 | 6.76 | 5.86 | 5.62 | 5.59 |

**D)**

| Media | pH (0h) | pH (10h) | pH (15h) | pH (20h) |
|---|---|---|---|---|
| LBG 0.1 | 6.77 | 4.04 | 3.72 | 3.61 |
| LBG 0.3 | 6.73 | 3.79 | 3.61 | 3.50 |

**Figure S6: A**) Kinetic plots showing the RFU values of the $P_{23}$_cymR variant when induced at different starting $OD_{600nm}$ (0.3 and 0.1) for 21 hours at 37 °C in LB or LBG media (n = 4, mean ± SD). **B**) Plots showing the kinetic growth (as $OD_{600nm}$) of the $P_{23}$_cymR variant (uninduced and induced) at different starting $OD_{600nm}$ (0.3 and 0.1) for 21 hours at 37 °C in LB or LBG media (n = 4, mean ± SD). “un” = uninduced condition, “in” = induced condition (5 μM of cumate). **C**) Table showing the pH values after 0, 10, 15, and 20 hours of incubation at 37 °C in LB medium and at starting $OD_{600nm}$ of 0.1 and 0.3 (n = 4). **D**) Table showing the pH values after 0, 10, 15, and 20 hours of incubation at 37 °C in LBG medium and at starting $OD_{600nm}$ of 0.1 and 0.3 (n = 4).

A)

| Media | Temp. | pH (0h) | pH (10h) | pH (15h) |
|---|---|---|---|---|
| MRS | 30 °C | 5.90 | 5.34 | 4.62 |
| MRS | 37 °C | 5.90 | 5.18 | 4.12 |

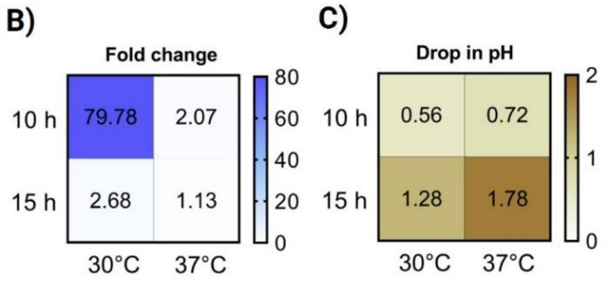


**Figure S7:** **A**) Table showing the pH values after 0, 10, and 15 hours of incubation at 30 and 37 °C in MRS medium at starting $OD_{600nm}$ of 0.01 (n = 4). **B**) Heatmap plot depicting the fold changes between the induced and uninduced conditions after 10 and 15 hours of growth at 30 and 37 °C in MRS medium and a starting $OD_{600nm}$ of 0.01. **C**) Heatmap depicting the drop in pH after 10 and 15 hours of growth at 30 and 37 °C in MRS medium and a starting $OD_{600nm}$ of 0.01. The pH values were deducted from the initial pH value (0 h) of each condition (30 and 37 °C).

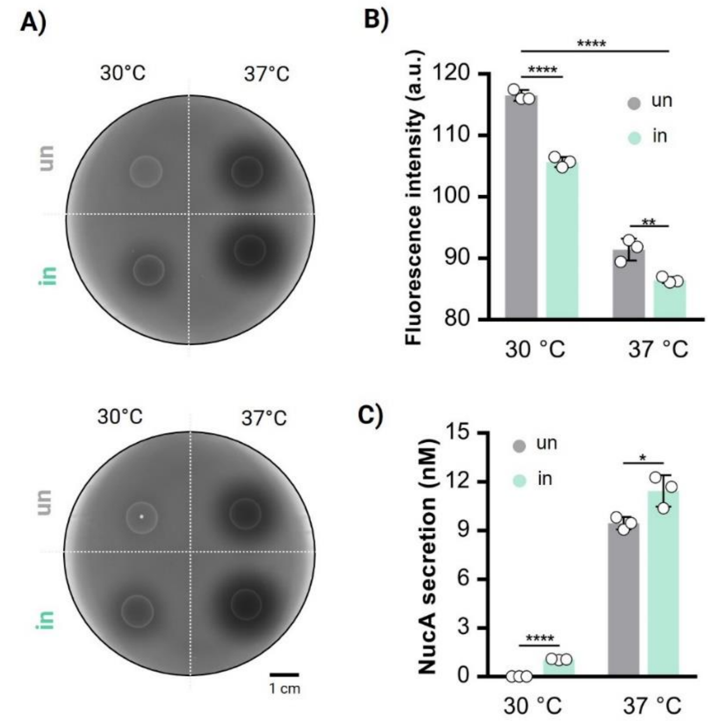


**Figure S8:** **A**) Amounts of NucA secreted (in nM) by $P_{23}$_cymR_NucA bacteria after growing in DNase medium for 24 hours at both 30 and 37°C and in the presence and absence of cumate (n = 3, mean ± SD). “un” = uninduced condition, “in” = induced condition (5 µM of cumate). **B**) DNase agar plates with 10 µL of spotted $P_{23}$_cymR_NucA bacteria, previously uninduced and induced for 8 hours in MRS medium at both 30 and 37°C, and incubated at 37°C for 40 hours. “un” = uninduced condition, “in” = induced condition (5 µM of cumate). **C**) Plot showing the levels of fluorescence intensity that were measured around the spotted area using ImageJ (n = 3, mean ± SD).

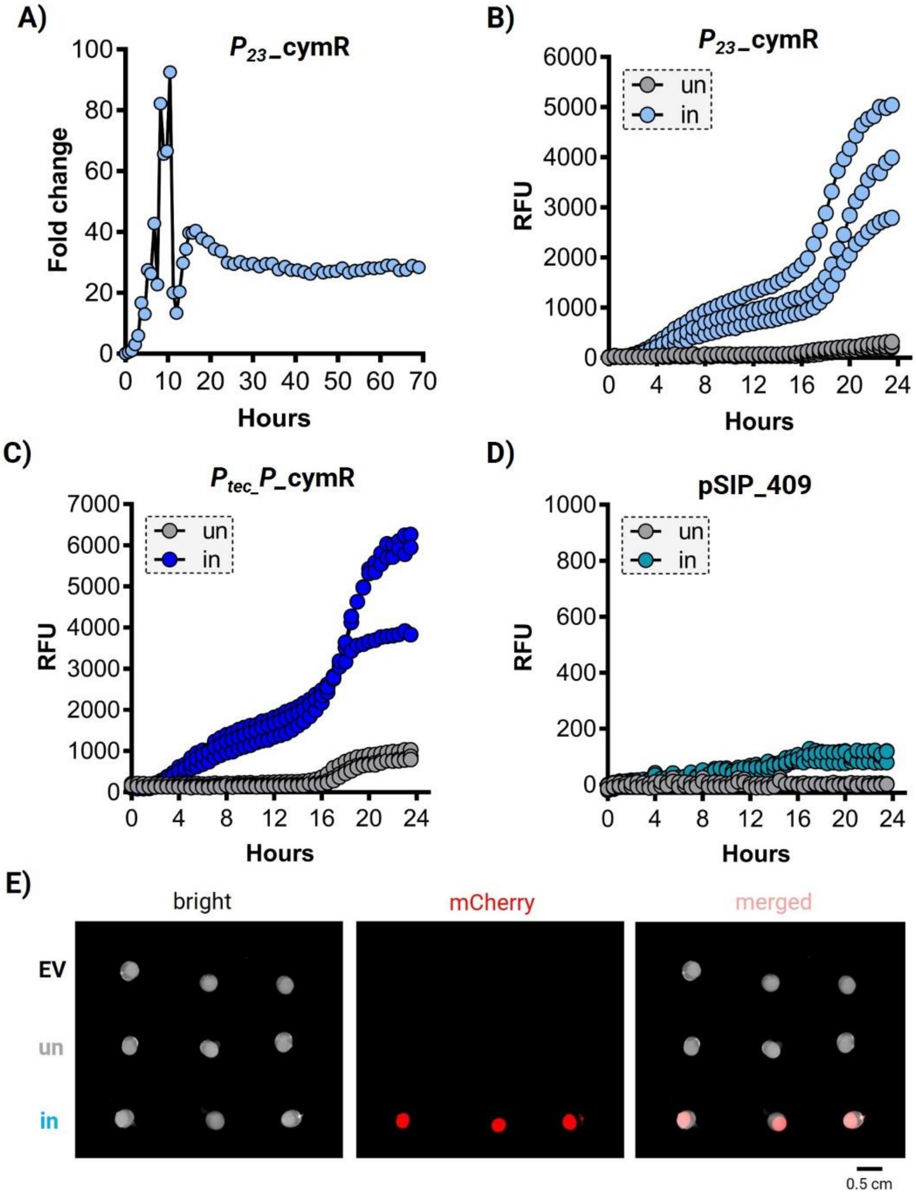


**Figure S9**: **A**) Fold changes for the induced state compared to the uninduced state for the $P_{23}$_cymR variant over 70 hours of growth at 37°C. **B**) RFU values of $P_{23}$_cymR-based PEARLs after induction with cumate for 24 hours (n = 3, mean ± SD). **C)** RFU values of $P_{tec}$_cymR_P-based PEARLs after induction with cumate for 24 hours (n = 3, mean ± SD). **D**) RFU for the mCherry-expressing pSIP409-based PEARLs when induced with IP673 for 24 hours (n = 3, mean ± SD). "un" = uninduced condition, "in" = induced condition (100 µM of cumate). **E**) BioRad Gel Documentation System images of EV-based PEARLs and the induced and uninduced conditions of $P_{23}$_cymR-based PEARLs after 24 hours of growth in DMEM cell culture medium and at 37°C (n = 3, mean ± SD). "un" = uninduced condition, "in" = induced condition (100 µM of cumate).

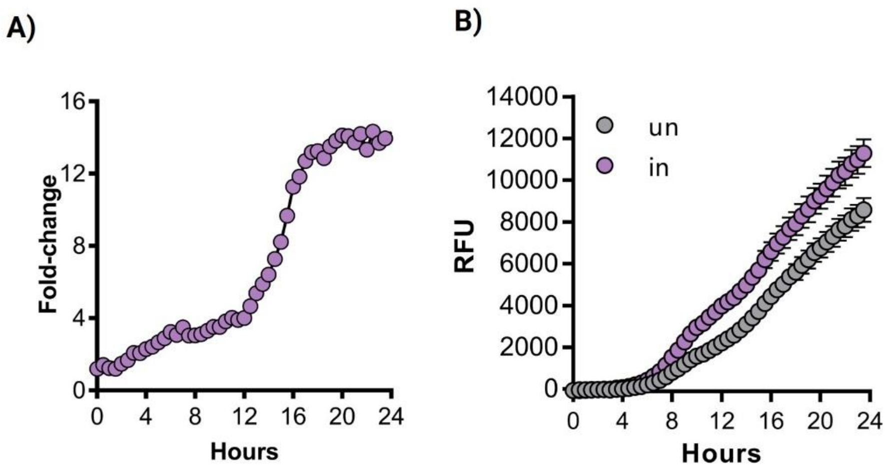


**Figure S10**: **A**) Fold-changes between the induced (with 100 µM of vanillate) and uninduced states of the $P_{23}$_vanR variant when encapsulated in PEARLs and grown at 37°C for 23 hours. **B)** RFU values corresponding to the induced and uninduced states of non-encapsulated $P_{23}$_vanR bacteria when grown at 37°C for 23 hours in culture (MRS medium) (n = 4, mean ± SD). “un” = uninduced condition, “in” = induced condition (100 µM of vanillate).

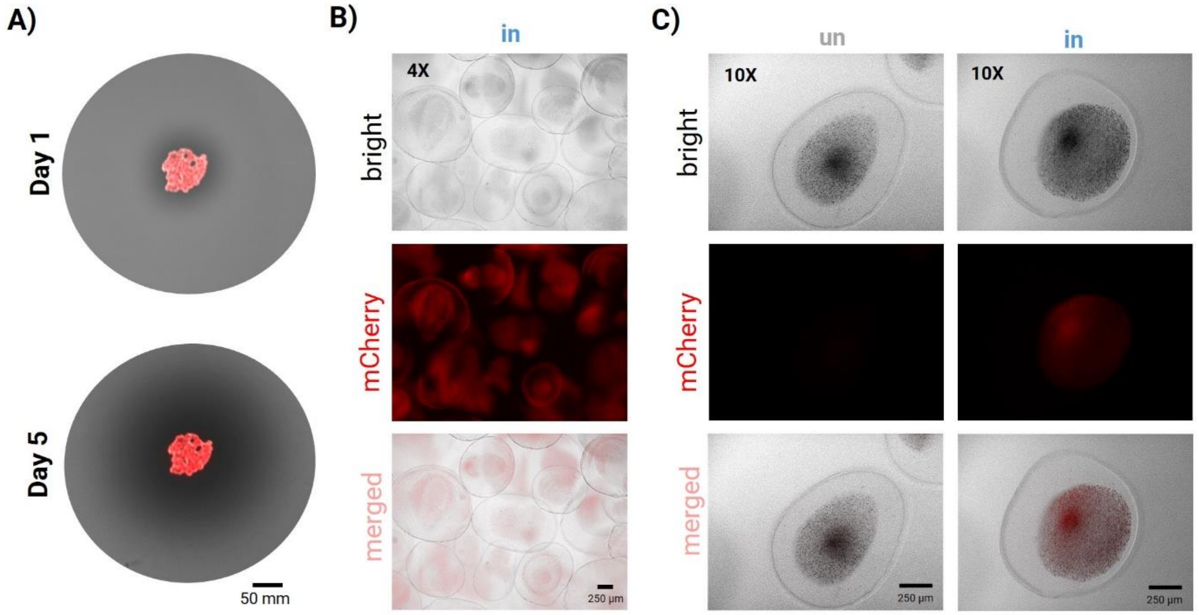


**Figure S11**: **A**) DNase agar plates with micro-PEARLs after 1 and 5 days of incubation at 37 °C. Images taken using the BioRad Gel Documentation System. **B**) Microscopy images at 4X of induced (in) micro-PEARLs after 24 hours of incubation at 37°C in a 12-well plate. “in” = induced condition (100 µM of cumate). **C**) Microscopy images at 10X of uninduced (un) and induced (in) micro-PEARLs after 24 hours of incubation at 37 ºC in a 12-well plate. “un” = uninduced condition, “in” = induced condition (100 µM of cumate).
